# The identification of a SARs-CoV2 S2 protein-derived peptide with super-antigen-like stimulatory properties on T-cells

**DOI:** 10.1101/2024.11.26.624714

**Authors:** Thai Hien Tu, Fatima Ezzahra Bennani, Nasser Masroori, Chen Liu, Atena Nemati, Nicholas Rozza, Amichai Meir Grunbaum, Richard Kremer, Catalin Milhalcioiu, Denis-Claude Roy, Christopher E. Rudd

## Abstract

Severe COVID-19 can trigger a cytokine storm, leading to acute respiratory distress syndrome (ARDS) with similarities to superantigen-induced toxic shock syndrome. An outstanding question is whether SARS-CoV-2 protein sequences can directly induce inflammatory responses. In this study, we identify a region in the SARS-CoV-2 S2 spike protein with sequence homology to bacterial super-antigens (termed P3). Computational modeling predicts P3 binding to sites on MHC class I/II and the TCR that partially overlap with sites for the binding of staphylococcal enterotoxins B and H. Like SEB and SEH peptides, P3 stimulated 25-40% of human CD4+ and CD8+ T cells, increasing IFN-γ and granzyme B production. viSNE and SPADE profiling identified overlapping and distinct IFN-γ and GZMB subsets. The super-antigenic properties of P3 were further evident by its selective expansion of T cells expressing specific TCR Vα and Vβ chain repertoires. *In vivo* experiments in mice revealed that the administration of P3 led to a significant upregulation of proinflammatory cytokines IL-1β, IL-6, and TNF-α. While the clinical significance of P3 in COVID-19 remains unclear, its homology to other mammalian proteins suggests a potential role for this peptide family in human inflammation and autoimmunity.

## Introduction

Coronavirus disease 2019 (COVID-19), caused by severe acute respiratory syndrome coronavirus 2 (SARS-CoV-2), continues to have severe health effects globally ^1, 2^. The virus enters host cells by the attachment of the spike (S) protein to the ACE2 receptors on cells ^3–5^. In this context, SAR-CoV-2 belongs to the beta coronavirus family of seven members that are known to infect humans^6^. Structurally, it is comprised of four structural elements that include the spike (S) glycoprotein, small envelope (E) glycoprotein, membrane (M) glycoprotein, and nucleocapsid (N) protein^6^. The S protein consists of two subunits, S1 and S2; the S1 subunit binds to target cell receptors at the receptor-binding domain (RBD), the S2 subunit mediates viral-cell fusion through its two-heptad repeat domain. SARS-CoV-2 vaccines are generally based on the use of the full-length S protein, the S1 subunit and RBD sequences^7^.

The more severe SARS-CoV-2 infections are often accompanied by an inflammatory cytokine storm^8–11^. Moreover, while mild cases involve antibody and T-cell responses, severe inflammatory disease is accompanied by reduced antibody responses^4, 12–17^ and an increase in pro-inflammatory cytokines such as interleukin-6 (IL-6)^18^. The effects of disease progression on the immune system ^9–19^ and the inflammatory response have now been well documented ^4, 12^. COVID-19 vaccines are now first-line preventative therapies inducing both B-cell and T-cell responses. In this context, T cells can recognize distinct conserved amino acid sequences across viral variants^20,21^. Severe disease is characterized by the preferential loss of TCF1+ stem-like progenitor T-cells^22^. While cytolytic CD8+ T cells can directly kill infected cells, CD4+ T cells can provide classic help for cellular and humoral responses^23^. In this sense, there is a correlation between the coordinated activation of B and T cell and SARS-CoV-2 clearance^12^, while greater T-cell receptor diversity is associated with milder COVID-19 symptoms^24–26^.

COVID-19’s inflammatory response including the COVID-19–associated multisystem inflammatory syndrome in children (MIS-C) shares some properties with toxic shock syndrome (TSS) ^27^. Clinical responses include a persistent fever and inflammatory responses within many organs. Severe infections are characterized by elevated levels of systemic inflammatory cytokines and weakened antiviral responses^27, 28^. In this context, TSS is often triggered by superantigens (SAgs) that can overstimulate the adaptive immune system. In contrast to antigen-induced T-cell responses, where only a tiny fraction of T-cells is activated, SAgs can activate up to 20% of the body’s T-cells^29–35^. Mechanistically, instead of binding to the antigenic peptide-binding site groove of MHCII, SAgs can bind to other regions as well as directly to the αβTCRs^36^. Examples include staphylococcal enterotoxins B (SEB) and H (SEH) which activate virus-specific CD8 T-cells directly, and indirectly via bystander effects ^37^. This hyper-activation can eliminate T-cells and/or induce a temporary state of hyperactivity followed by unresponsiveness^38,39^. In this context, the expression of human endogenous retrovirus HERV-K18 superantigen has been connected linked to juvenile rheumatoid arthritis ^40^, while V-beta-restricted T cell adherence to endothelial cells is linked to vascular injury ^41^. Other endogenous superantigens also exist in humans ^42,43^.

Although the existence of SARs-CoV-2 derived peptides such as T_678_NSPRRAR_685_ with SAg binding properties has been reported^44^, peptides with an ability to induce T-cell activation have yet to be identified. In this study, by comparing sequences and homologies to known bacterial super-antigens, we identified an additional peptide region in the S2 protein that can induce broad low-level activation of CD4 and CD8+ T-cells. Virtual docking experiments confirmed the peptides P3 and related P3b bind to MHC class I/II and TCR sites. Moreover, the administration of P3 resulted in the *in vivo* generation of IL-6, IL-1b, and TNF-α cytokines within mice, along with an amplification of specific subsets of TRAV and TRBV in T-cells. Lastly, it is noteworthy that P3 exhibits homology with endogenous proteins present in mammalian cells, thereby raising the possibility of related endogenous peptides exist that could stimulate autoimmunity and inflammation.

## Results

### Peptide from SARs CoV2 S2 spike protein exhibits a significant binding interaction to the TCR and MHC class I and II complexes

Given that SARS CoV2 infections can involve inflammatory response, we asked whether there might be regions of the SARs spike protein that stimulate T-cells in a manner similar to super-antigens (SAgs). For this, we carried out initial comparisons between the sequence of the Spike and superantigens that induce septic shock syndrome. In addition to T_678_NSPRRAR_685_^44^, we identified another region in the S2 spike peptide (D_1139_PLQFPELDSFKEEL_1152_, termed P3) with homology with regions in enterotoxins SEB and SEH (Supplementary Table 1). We also analyzed the homology of P3 to other coronaviruses and in SARS-CoV-2 variants (α, β, γ, δ, and Omicron). As shown in Supplementary Table 2, P3 showed the most homology with SARS-CoV-2 (WT) (66.67%) and to a lower extent with other seasonal coronaviruses (OC43, NL63, HKU1 and 229E) (around 30-45%). Further, P3 is conserved with the same percent similarity (66.67%) to five SARS-CoV-2 variants (Supplementary Table 3).

We next conducted molecular docking simulations to assess the potential binding of peptide P3 to the HLA class I/II-TCR complexes similar to SEB or SEH (Figs. 1 and 2). Importantly, peptide 3 showed molecular docking with HLA-I-TCR protein (PDB: 3PWP) (Fig. 1a-b). This showed that both P3 and a derived SEH peptide bound to the interface connecting HLA-I to TCR. Nine amino acids of P3 created intermolecular interactions with the complex HLA-I-TCR. We observed P3 H-bond interactions with residues Thr73, Thr80, Gln96, Arg97, Trp133, Thr142, and Lys146 of HLA-I and residues Gln105, Phe106 of TCRα besides Glu30, Leu98, Ala99, Gly101 of TCRβ (Fig. 1c). A salt bridge interaction was noted on residue Lys146 of HLA-I and Glu30 of TCR. The comparison of the binding of P3 and SEH to HLA-I-TCR reveal that they both make H-bonds with residues Thr73, Thr142 of HLA-I and on residues Gln105, Leu98, Ala99 of the TCR (Fig. 1c-e).

**Fig. 1.**
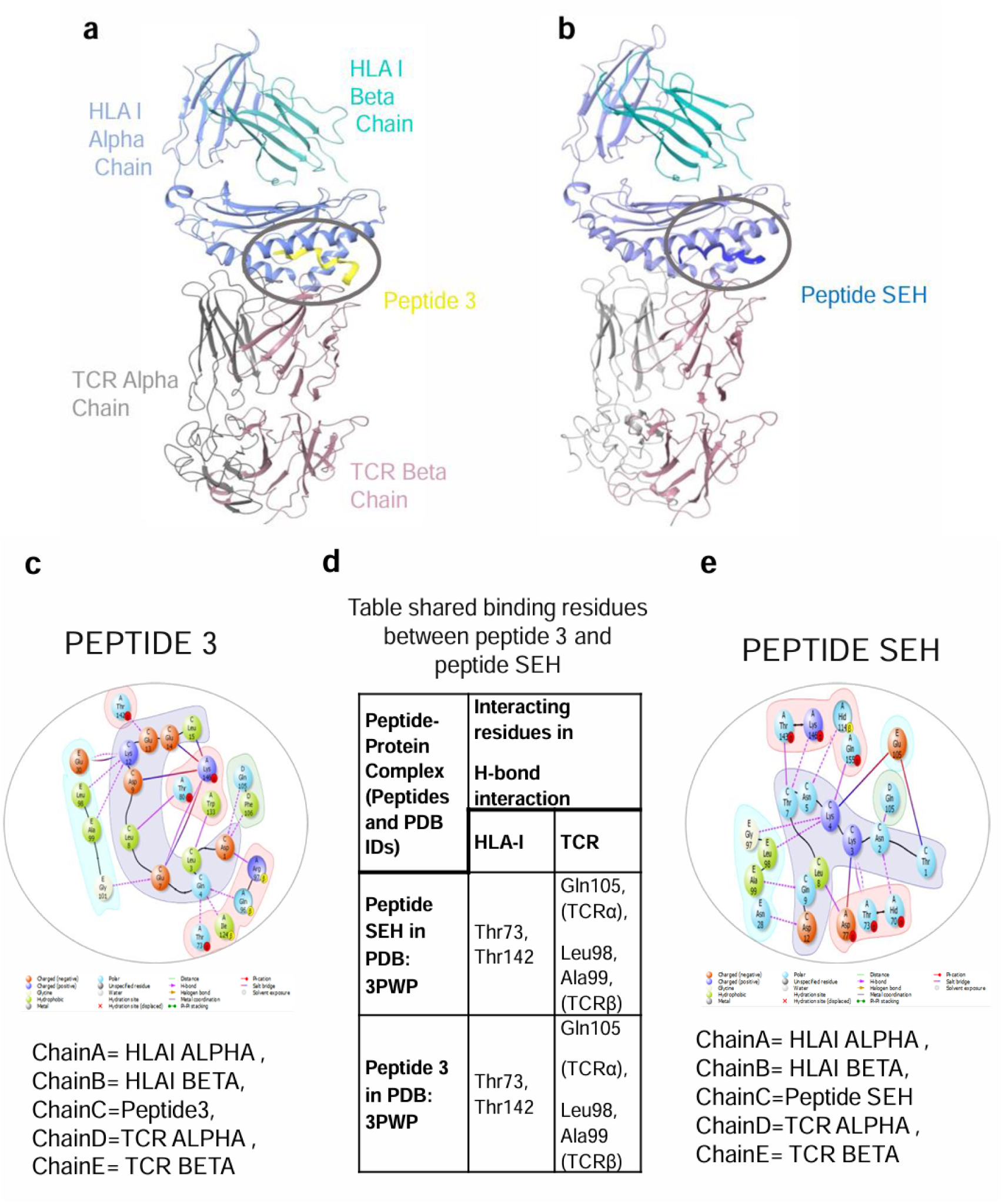
Overview of three- and two-dimensional structure of the complex docking TCR– HLA-1 peptides complex (PDB ID 3PWP). **a** Complex docking of peptide 3 with TCR and human class MHC (HLA) I antigen displayed in cartoon style for HLA-I_TCR and peptide 3 is yellow ribbon, showing binding at the interface connecting TCR to HLA-I. Peptide 3 makes 14 H-Bond interactions and 6 salt-bridge interactions. **b** Complex docking of peptide SEH with TCR and human class MHC (HLA) antigen displayed in cartoon style for HLA-I_TCR and peptide 6 is dark blue ribbon, showing binding at the interface connecting TCR to HLA-I. **c** 2D diagram of interaction of peptide 3 and SEH with MHC-class 1 and TCR residues. Left panel: colored diagram showing binding residues for P3; Middle table: Table showing the H bond binding to residues for peptide 3 and peptide SEH binding to the complex HLA-I. Right panel: colored diagram showing binding residues for peptide SEH. SEH makes 11 H-Bond interaction and 3 salt-bridge interactions.

**Fig. 2.**
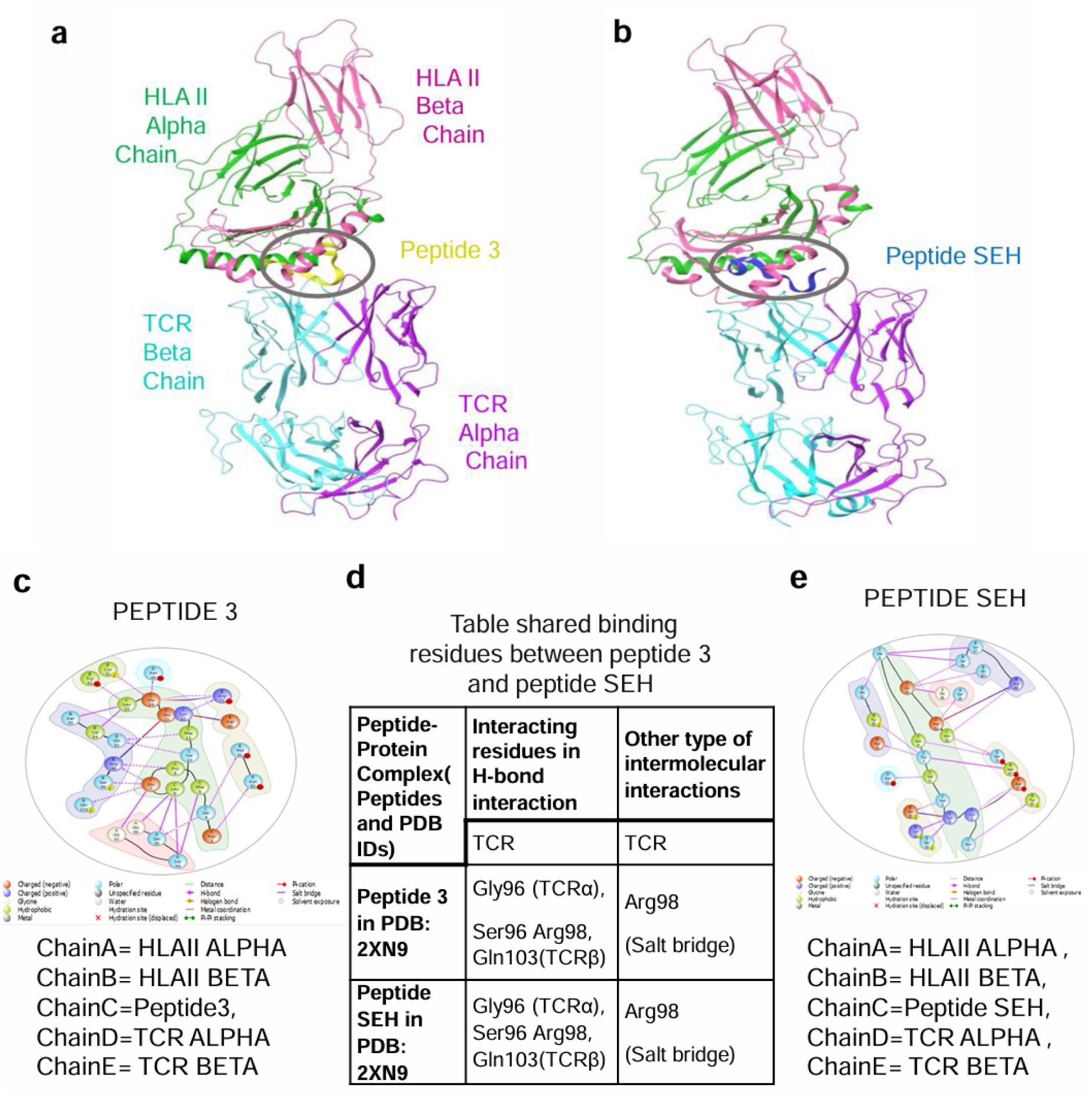
Overview of three- and two-dimensional structure of the complex docking TCR– HLA-II peptides complex (PDB ID 2XN9). **a** Complex docking of peptide 3 with TCR and human class MHC (HLA) II antigen displayed in cartoon style for HLA-II_TCR and peptide 3 is yellow ribbon, showing binding at the interface connecting TCR to HLA-II. Peptide 3 makes 20 H-Bond interactions and 3 salt-bridge interactions. **b** Complex docking of peptide SEH with TCR and human class MHC (HLA) II antigen displayed in cartoon style for HLA-II_TCR and peptide SEH is dark blue ribbon, showing binding at the interface connecting TCR to HLA-II. SEH make 11 H-Bond interaction and 3 salt-bridge interactions. **c** 2D diagram of interaction of peptide 3 and SEH with MHC-class II and TCR residues. Left panel: colored diagram showing binding residues for P3; Middle table: Table showing the H bond binding to residues for peptide 3 and peptide SEH binding to the complex HLA-II. Both P3 and SEH make H-bond and salt bridge on Arg98. Right panel: colored diagram showing binding residues for SEH.

Molecular docking analysis also showed an interaction profile for P3 with the HLA class II-TCR protein (PDB: 2XN9) complex (Fig. 2). The HLA II-TCR protein crystal structure was retrieved from the ternary protein complex of staphylococcal enterotoxin H and was docked with the P3 and the SEH peptide for comparison. The best generated complexes showed that both peptides are binding in the same region as SEH (Fig. 2a-b). This revealed multiple non-covalent interactions for peptide 3, such as H-bond interactions with both the proteins (Fig. 2c). This included amino acids residues Cys30, Trp61, Arg71, His81, Asn82 and Asn69 on HLA-II which participated in H-bond interactions with Peptide 3. In addition, several residues of TCR protein such as Gly93, Ser94, Gln95, Gly96 (TCRα), Gln52, Val54, Asn55, Ser96, Arg98, and Gln103 (TCRβ) establish H-bonds with P3. In addition, salt bridge interactions were observed between P3 and residues Asp66, Arg71 of HLAII, and residue Arg98 of TCR complex. Mostly the penta-(QFPEL) and hexa-peptide (SFKEEL) unit of P3 mediated the H-bond and salt bridge interaction with the HLAII-TCR complex. After comparison of 2D diagram of both peptide3 and SEH, we observed that they shared 4 H-bond on the same residues that are Gly96(TCRα) and Ser96, Arg98, Gln103(TCRβ) (Fig. 2c-e). Although the interaction profiles of P3 with 2XN9 and 4C56 were slightly different, the orientation of the binding to the interaction interface region remained similar to the binding of staphylococcal enterotoxin H or B. P3 binding also partially overlapped with the binding of full-length SEB to the MHC-TCR complex (Supplementary Fig. 1). These data show that P3 peptide can bind to an interactive region between HLA class 1 and 2 and the TCR sequences in a manner like SEB, SEH peptides and full-length SEB.

In our initial search of Spike sequences, we first identified this unusual variant with the sequence DPLQFPELDSFKEEL (P3); however, later searches showed the dominant sequence DPLQPELDSFKEEL (we have now termed P3b) which differed by the presence of a single phenylalanine (F) (Supplementary Table 1, upper panel). Using CABSDOCK and HPEPdock, we also performed Blind and Local Docking and showed that they bind to a similar region between MHC class I and II and the TCR. P3 and P3b showed differences in binding to specific residues in class I antigens where P3b made numerous H-bond interactions with HLA1 and a salt bridge with TCR (Supplementary Fig. 2a-e). In the case of class II, P3 and P3b binding was observed to a similar region involving similar and distinct binding sites. Both peptides shared H-bond interactions with TCRβ residues Gln52, Val54, Asn55, Arg98 (Supplementary Fig. 3a-e). In sum, while differences were noted in certain interactive residues, both P3 and P3b bound to similar regions, and contacted both the TCR and MHC antigens.

### Peptide 3 induces T-cell proliferation

Given the potential binding to TCR and MHC, we next determined whether P3 could activate human peripheral blood T-cells from healthy donors (Fig. 3). As controls, we included the use of peptides from superantigen SEB and SEH (n=4). T-cells labeled with Tag-it violet cell tracking dye were incubated with the various peptides for 6 days followed by an analysis of cell proliferation by flow cytometry (see *Methods)*. The peptides were titrated to obtain the optimal concentration for the activation of T-cells (Supplementary Fig. 5). Firstly, the P3 peptide stimulated some 30-40% of T-cells when compared to the media control (Fig. 3a). P3 peptide also stimulated cells within the CD4+ (Fig. 3b) and CD8+ subset (Fig. 3c). By contrast, the SEB and SEH peptides also activated 20-25% of CD4+ T-cells but only weakly activated CD8+ T-cells (i.e., 5%).

**Fig. 3.**
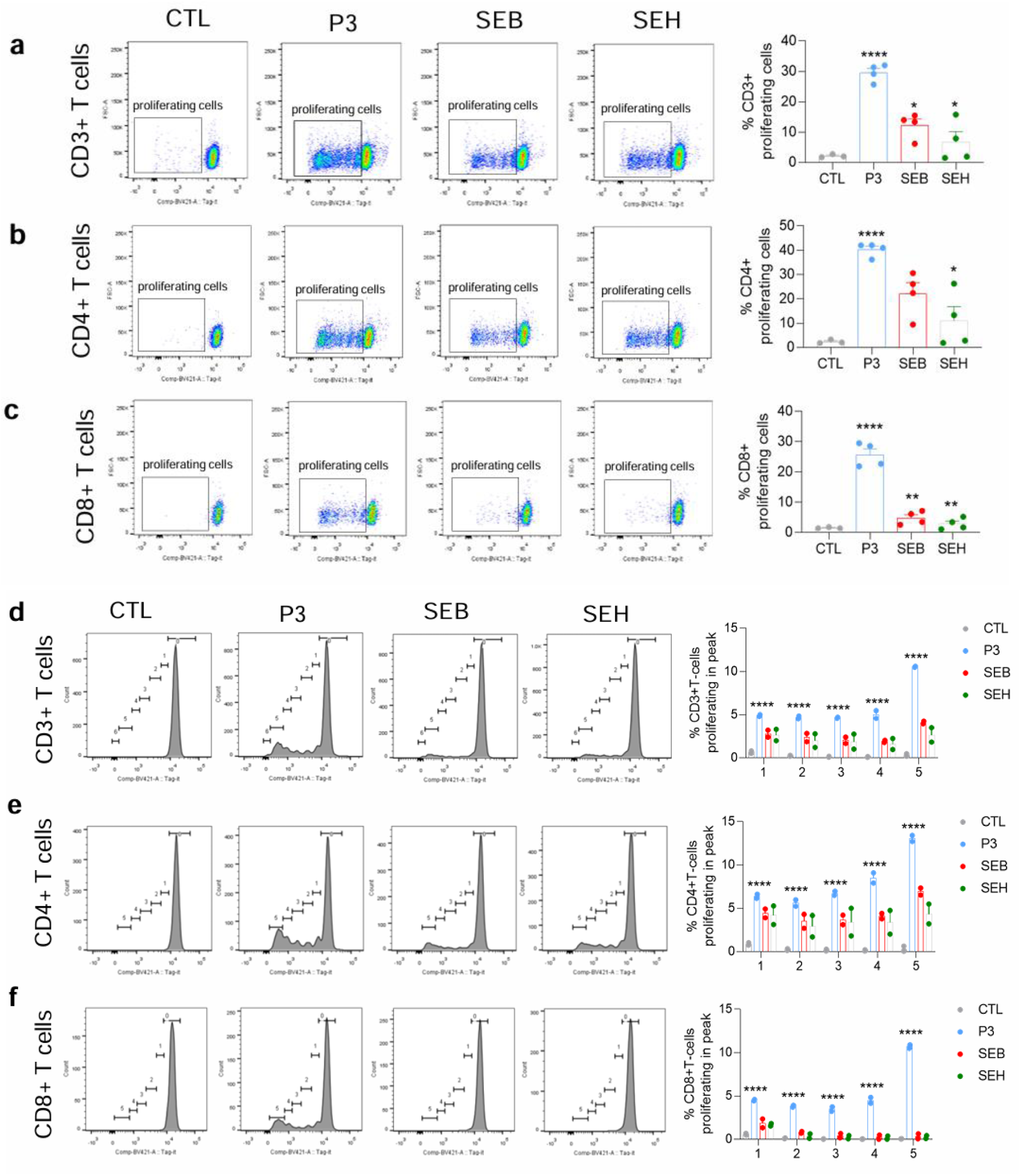
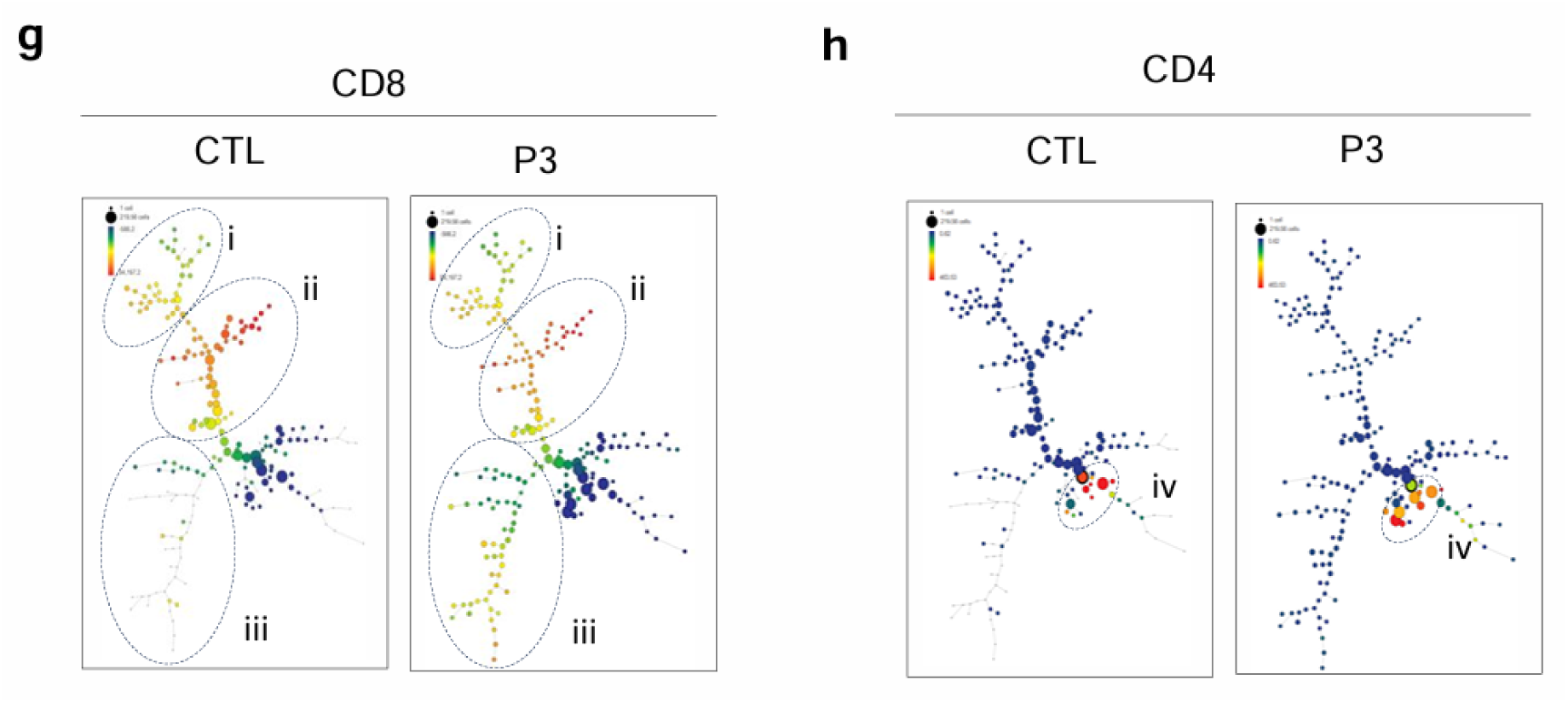
P3 peptide spike protein peptide enhances human T-cell proliferation. PBMCs cells from 4 donors were stained with Tag-it before incubating with peptide 3, SEB and SEH; media only without adding peptide was used as negative control group. The percentage of proliferating cells were measured with Tag-it proliferation assay after a 6 days incubation. **a-c** Left panel: Representative flow cytometry plots displaying gating for proliferating cells in T-cells (**a**), CD4+ (**b**) and CD8+ (**c**) T-cells. Proliferating cells are gated (left gates). Right panel: Histogram data show the percentage of proliferating cells. Data are presented as Mean ± SEM from 4 human PBMCs samples, significant at p ≤ 0.05 *, p ≤ 0.01 ** and p ≤ 0.001 ****. **d-f** Left panel: The number of divisions was determined in proliferating T-cells (**d**), CD4+ (**e**) and CD8+ (**f**) T-cells. The gate “0” capturing the Tag-it undivided cells on the right and Tag-it divided cells on the left is identified with 5 divisions. The gate labeled as “0,1,2,3,4,5” indicate the numbers of cell division. Right panel: Histogram showing the percentage of proliferating cells in 5 divisions; CD3+ T-cells, CD4+ T-cells, CD8+ T-cells. Data are presented as Mean ± SEM from 2 human PBMCs samples, significant at p ≤ 0.001 ****. **g-h** SPADE analysis shows the increasing CD8 and CD4 population in CD3+ T-cells, with peptide 3 treatment. The fluorescence intensity of different markers for each node are represented by color (green to red), while the size of node represents the number of cells. Peptide increases the node in tree grouping iii (CD8+ cells) and tree grouping iv (CD4+ cells). The SPADE analysis was run with concatenated samples from 4 human PBMCs samples.

We next monitored the number of divisions in the proliferating cell population (Fig. 3d-f). The P3, SEB, and SEH peptides induced the same number of divisions (5 divisions). After 6 days of culture, P3-treated T-cells progressed through five divisions, with 4-6% of cells having undergone 4 divisions and 10% with a 5^th^ division (Fig. 3d). SEB or SEH peptide-treated cells showed some 2-3% with 1,2,3,4 divisions and 4% with a 5^th^ division (Fig. 3d). A similar pattern of cell division was seen in CD4+ T-cells (Fig. 3e). It is noteworthy that P3 induced 3-4% of CD8+ T-cells in division 1,2,3,4 and 10% in division 5, while SEB and SEH peptides induce less than 1% cells (Fig. 3f). The addition of the P3b peptide to cultures of mouse T-cells also induced their activation as shown by the expansion of a subset of CD4 and CD8 T-cells by 4-5 cell divisions (Supplemental Fig. 6). Furthermore, the SPADE analysis well defined CD8 and CD4 population in P3-treated cells as compared with CTL, three tree groupings (i-iii) are CD8+ cells and group iv is CD4+ cells (Fig. 3g-h). We found the incubation with P3 caused an increase in the numbers and size nodes in group iii (Fig. 3g) and iv (Fig. 3h) which denoted increased numbers of CD8+ and CD4+ cell population.

### Peptide 3 induces activation and cell cycling markers

With a focus on P3, we next assessed whether the peptide could induce activation markers CD69 and Ki67. CD69 is an activation antigen induced by the TCR ligation ^45^. Firstly, we defined cellular populations using high-dimensional spanning-tree progression analysis for density-normalized events (SPADE) ^46^. SPADE revealed an extensive heterogeneity within the CD4 and CD8 subsets (>80 nodes of distinct cells) with multiple tree branches depicting different cell groupings within the population. Within each pattern, we focused on 7 groupings of tree clusters and their nodes (i-vii). Most of the cells in the control CD4+ T-cells (CTL) were found in four tree groupings i-iv (Fig. 4a). Most nodes either showed a low level (light green) or no expression (dark green). With P3, there was a reduction in the size of some nodes in tree grouping iii and iv accompanied by the increased presence of cells in grouping v, vi, and vii, and grouping v has high levels of CD69 expression (circled, orange, and red).

**Fig. 4.**
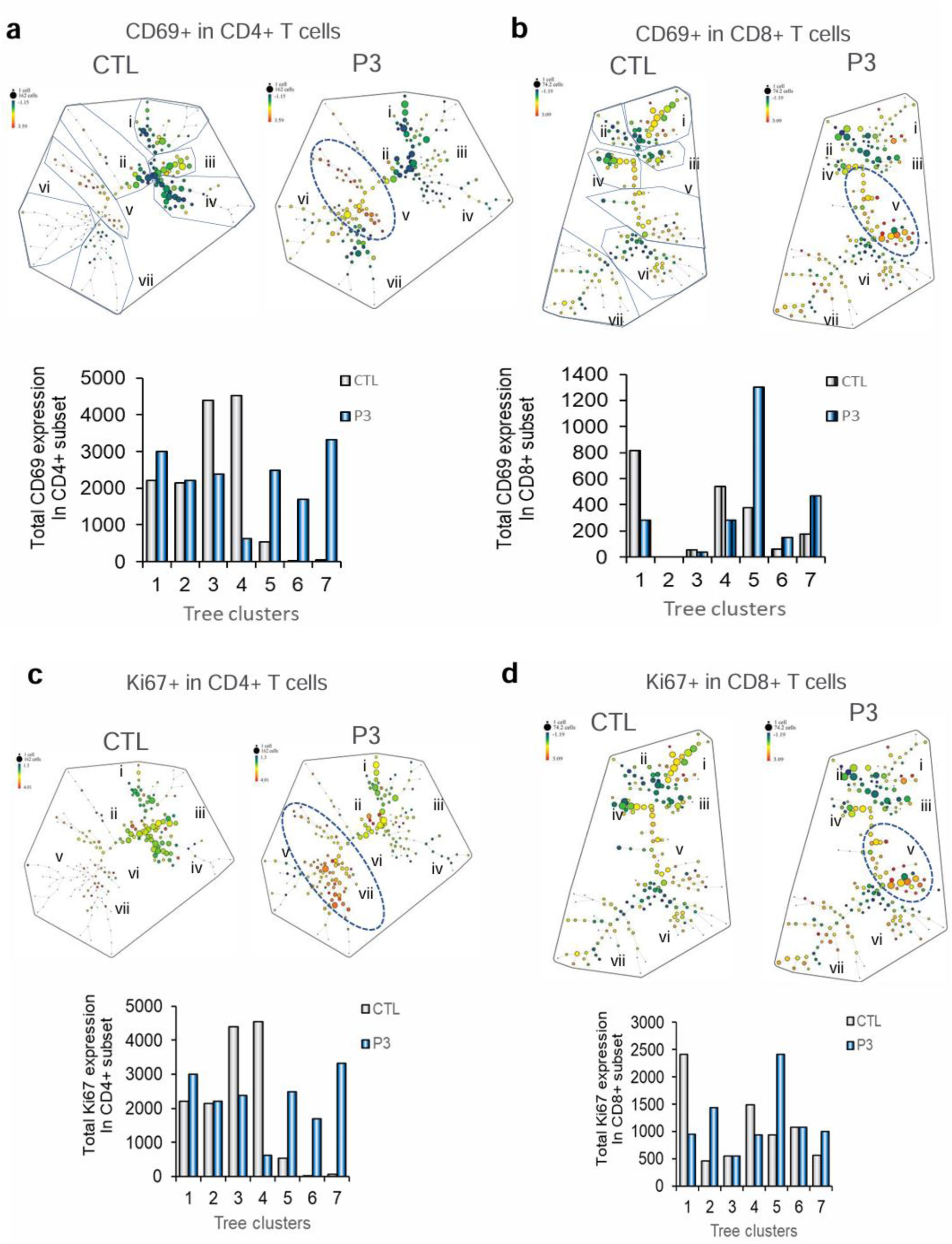
P3 spike peptide activates CD4+ and CD8+ human T-cells. Human PBMCs (n=4) were incubated with peptide 3, SEB and SEH for 6 days, followed by an analysis of activation markers by flow cytometry. **a** CD69 expression on CD4+ T-cells. SPADE patterns showing the increase of CD69 expression in CD4+ T-cells with peptide 3 treatment. The fluorescence intensity of different markers for each node are represented by color (green to red), while the size of node represents the number of cells. Histogram showing the total MFI level of CD69 expression in each tree cluster in CD4+ T-cells. Data is presented as Mean value of 4 human PBMCs concatenated sample. **b** SAPE analysis for CD69 expression on CD8+ T-cells. Histogram showing the total MFI level of CD69 expression in each tree cluster in CD8+ T-cells. Data is presented as Mean value from 4 human PBMCs concatenated sample. **c** SAPE analysis showed Ki67 expression on CD4+ T-cells. Histogram showing the total MFI level of Ki67 expression in each tree cluster in CD4+ T-cells. Data is presented as Mean value from 4 human PBMCs concatenated sample. **d** Ki67 expression on CD8+ T-cells by SPADE analysis and Histogram showing the total MFI level of Ki67 expression in each tree cluster in CD8+ T-cell. Data is presented as Mean value of 4 human PBMCs concatenated sample. The analysis incorporates concatenated data from 4 human samples in each condition. Equal numbers of cells in each condition were analyzed.

Next, we quantified the level of CD69 expression where P3 induced higher expression of CD69+ cells in the CD4 T-cells subset relative to resting controls (Fig. 4a, lower panels). Similarly, there was a complex array of different nodes within the CD8+ SPADE pattern, i-vii (Fig. 4b). The transition from resting to P3 activated cells involved the loss of cells in tree grouping i followed by the appearance of new cells in grouping v. Grouping v showed an increase in the expression of CD69 (see red and orange (circled at right panel). P3 therefore increased the expression of CD69 in a subset of CD4 and CD8+ T-cells consistent with the stimulation of a subpopulation of cells.

We next examined the cell cycling by measuring the expression of marker Ki67 (Fig. 4c-d). The nuclear protein Ki67 is expressed by cycling and recently divided cells, but not by naïve or resting cells^47^. P3 induced Ki67 on 30-35% of CD4+ T-cells and 25-35% of CD8+ T-cells (Supplementary Fig. 7a). There was an increase in the MFI for expression with P3, SEB, and SEH peptides; and the increase by incubation P3 was greater than with SEB and SEH peptides (Supplementary Fig. 7b). Indeed, in the case of CD4+ T-cells, SPADE analysis showed low levels of Ki67 expression in four tree groupings (i-iv) (Fig. 4c). Incubation with P3 caused an increase in the numbers and size of nodes in grouping v, vi, and vii which concurrently showed an increase in the expression of Ki67 (circled) (Fig. 4c). In the case of CD8+ T-cells, most cells were found in the nodes in three tree groupings i-iii. P3 induced a reduction in tree groupings i that was accompanied by an increase in the presence and expression of Ki67 in tree grouping v (orange and red; circled at Fig. 4d). These data showed that P3 increased the cell cycling and activation markers in the same population of subsets of both CD4 and CD8 T-cells. This theme was also seen by viSNE analysis. The cell densiometric profile for control cells showed several clusters which underwent substantive changes in distribution with P3, SEB, and SEH activation in the case of CD4+ T-cells (Supplementary Fig. 7c-d). P3 had the most marked effect in inducing a major regrouping of cells. This was accompanied by an increase in CD69 and Ki67. The expressions of CD69 and Ki67 appeared on different subsets of cells. In the case of CD8+ T-cells, P3 lead to the complete loss of cluster 1 accompanied by the appearance of the expression of Ki67 on cluster 3. SEB and SEH had only minor effect on the distribution of cells, consistent with the weak proliferation data and weak induction of Ki67 expression (Supplementary Fig. 7b). Interestingly, the expression of CD69 was higher on SEB and SEH activated cells. As in the case of CD4+ T-cells, the expressions of CD69 and Ki67 appeared on different subsets of cells.

### Peptide 3 induces the expression of effector molecules

We next assessed whether P3 could induce the expression of effector molecules such as the cytokine IFN-γ and GZMB. P3 induced a marked increase in IFN-γ expression on 34% of CD4+ cells (Fig. 5a). This percentage was much higher than seen with SEB or SEH and as such was the marker that most distinguished the effects of P3 on T-cell activation. SPADE analysis further showed P3 increase in expression of IFN-γ at node groupings v, vi and vii (red color; circled at right upper panel).

**Fig. 5.**
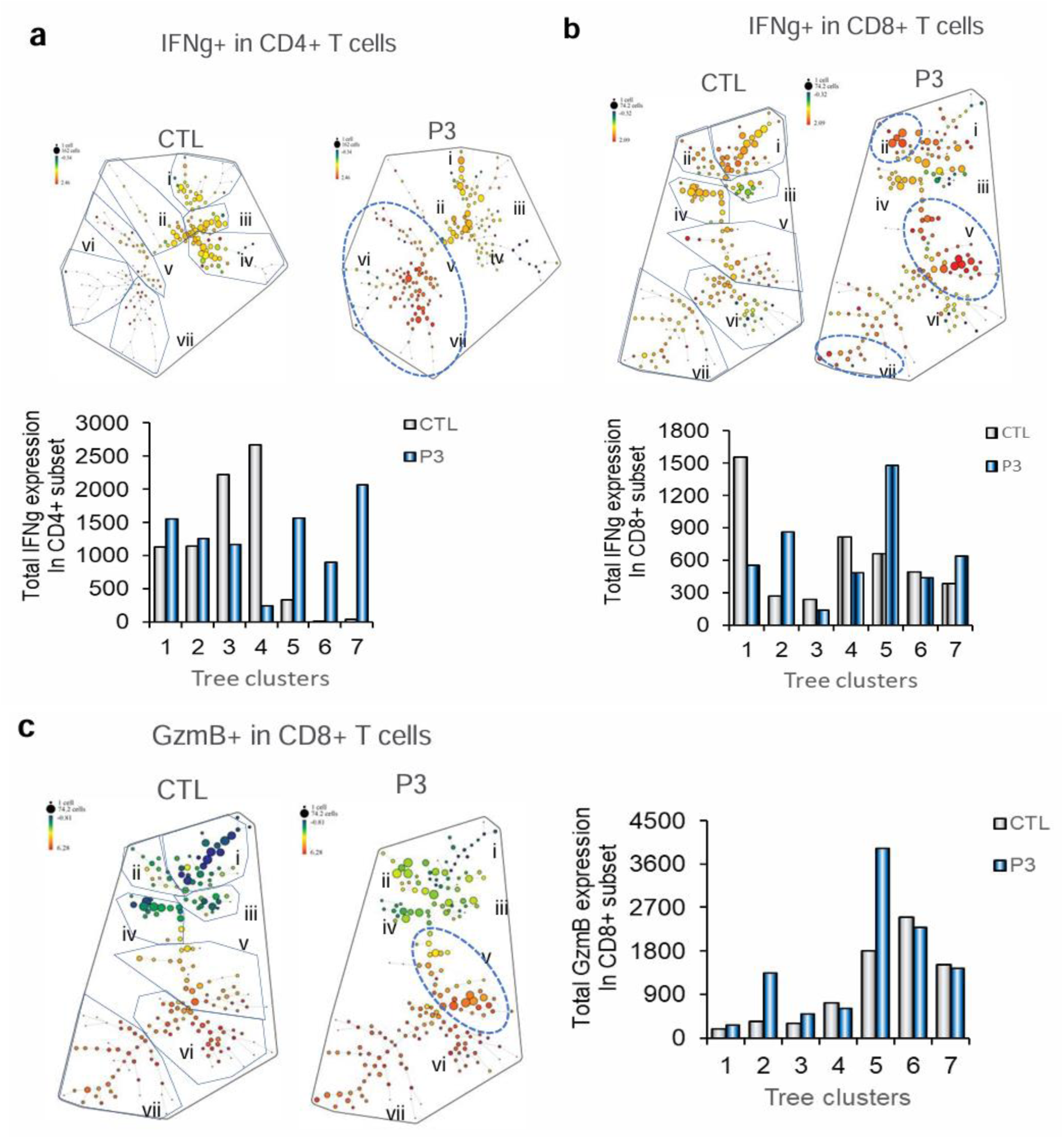
P3 spike peptide induces the expression of effector molecules in CD4+ and CD8+ human T-cells. Human PBMCs (n=4) were incubated with peptide 3, SEB and SEH for 6 days, followed by an analysis of activation markers by flow cytometry. **a** IFNg expression on CD4+ T-cells. SPADE patterns showing the increase of IFNg expression in CD4+ T-cells with peptide 3 treatment. Histogram showing the expression levels of IFNg+ cells in each tree cluster in CD4+ population due to incubation with the peptide 3. Data is presented as Mean value of 4 human PBMCs concatenated sample. **b** IFNγ expression on CD8+ T-cells. SPADE patterns showing the increase of IFNg expression in CD8+ T-cells with peptide 3 treatment. Histogram showing the expression levels of IFNg+ cells in each tree cluster in CD8+ population due to incubation with the peptide 3. Data is presented as Mean value of 4 human PBMCs concatenated sample. **c** Percent of granzyme B (GZMB) expression on CD8+ T-cells. Description same as b. The analysis incorporates concatenated data from 4 human samples in each condition. Equal numbers of cells in each condition were analyzed.

P3 induced expression in 18-20% of CD8+ cells (Fig. 5b). SPADE showed the control samples has expression of IFN-γ at low levels in three tree grouping i-iii, while P3 caused an increase in expression of IFN-γ at nodes grouping ii, v and vii and a loss of cells in tree grouping i. These data confirmed that expression of IFN-γ was induced on a subset of CD4 and CD8+ human T-cells.

Lastly, we also observed an increase in GZMB on CD8+ T-cells (Fig. 5c). The increase occurred from 30% on control cells to 40% on P3 activated cells. SPADE confirmed the increase in tree grouping v (circled). A comparison of the SPADE patterns for CD69, Ki67, IFN-γ and GZMB expression showed in increase in CD8+ cells within tree grouping v. A subset of cells therefore was activated to undergo cell division and express effectors IFN-γ and GZMB. At the same time, other subsets such as subset ii showed an increase in the expression of IFN-γ but not GZMB. These different subsets would therefore be expected to mediate different effect functions in response to P3.

### P3 peptide promotes inflammatory cytokines production in T-cells

Given that P3 can act to stimulate T-cells similar to SEB and SHE peptide, we next assessed its ability to induce pro-inflammatory cytokines (Fig. 6). In severe diseases, lymphopenia is correlated with high levels of proinflammatory cytokines ^9,10, 18, 48, 49^. To this end, we confirmed increased cytokine levels in COVID-19 patients’ serum where IL-6, IL-8 and chemokine MCP1 correlated with disease severity as compared to healthy controls (Supplementary Fig. 8). High levels of IL 6 are associated with a poor prognosis in COVID-19^50,51^. For this, peripheral T-cells from healthy patients were cultured with the P3 peptide for 6 days followed by an assessment of IL-6 transcription on isolated T-cells by RT-PCR. As controls, we also measured the transcription of TNF-α and IL-1β. P3 induced a marked increase in the transcription of IL-6 by more than 50-fold and IL-1β by two-fold. No change in TNF-α levels was seen.

**Fig. 6.**
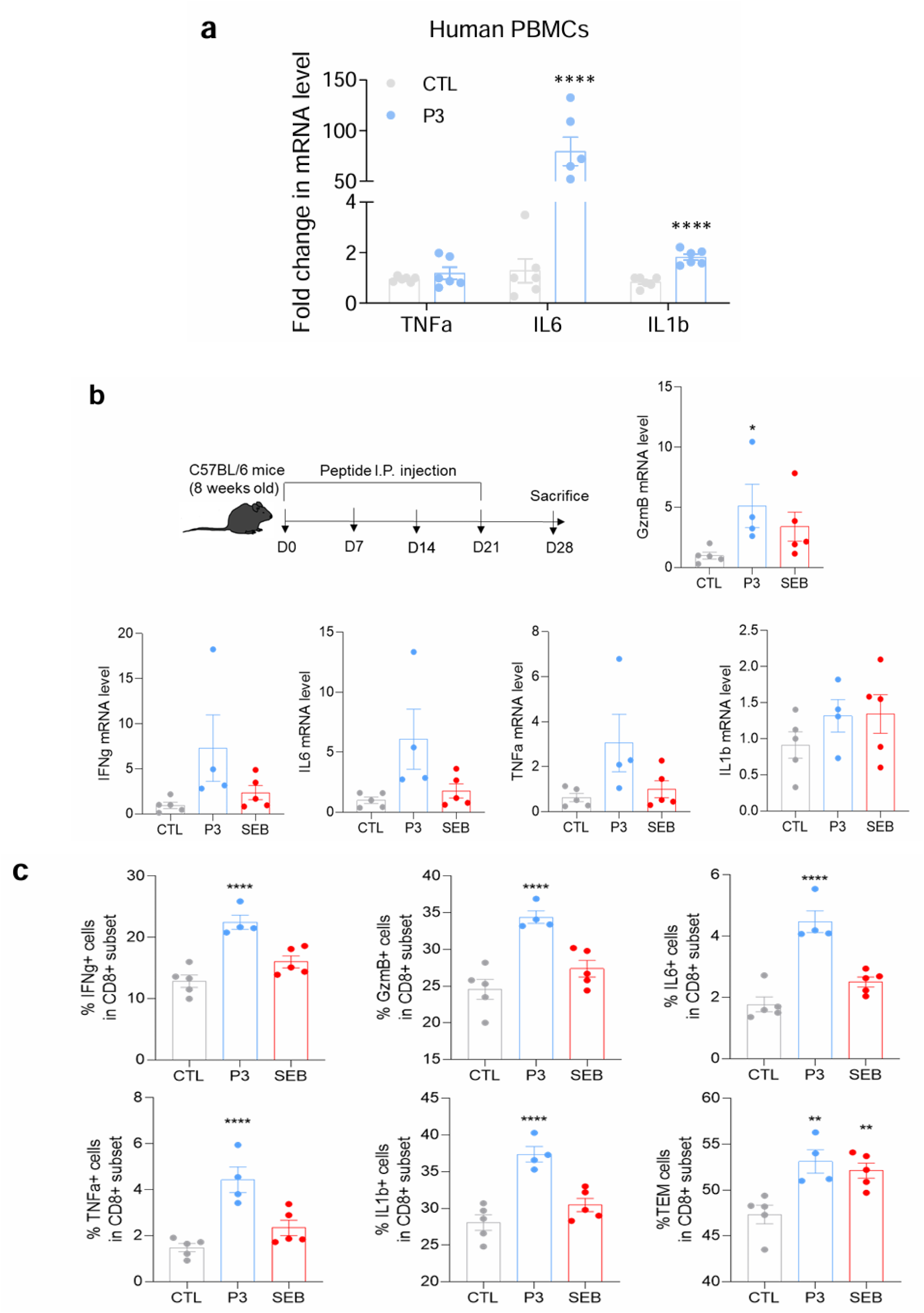
P3 induces proinflammatory cytokines in CD8+ human T-cells and *in vivo* in mice. **a** Human PBMCs were incubated with peptide 3 for 6 days, and CD8+ T cells were isolated (n=3). Total RNA was extracted, followed by RT-PCR analysis. Expression of inflammatory cytokines IL6 and IL1b in peptide 3-treated cells. Cytokines mRNA levels were measured by real-time PCR, then normalized to the level of human 18S mRNA. Data are presented with fold change different as compared with control sample. Data represents the mean ± SEM of technical duplicate. p ≤ 0.05 *, p ≤ 0.01 **. **b** Schematic representation of in vivo experiment. C57BL/6 mice were injected with P3 or SEB once a week for 3 weeks. At the end of experiment, spleen was harvested after 1 week of last injection for investigation. The mRNA levels of cytokines (IFNg, TNFa, IL6, GzmB, IL1b) in spleen of experimental mice were measured by real-time PCR, then normalized to the level of house keeping gene b-actin. Data are presented fold change different as compared with control sample. The significant differences were compared between P3-treated mice or SEB-treated mice with control group (no peptide treatment). **c** Secretion protein levels of IFNg, GzmB, IL1b, IL6, TNFa cytokines in splenocytes of mice immunized with peptide 3 or SEB. The levels of cytokines were determined by using flow cytometry, showing the percentage of CD8+ T cell expressing each cytokine. Data was analyzed and presented as mean ± SEM from 5 mice in CTL group, n=4 mice in P3-injected group and n=5 in SEB-injected group; significant at p ≤ 0.01 ** and p ≤ 0.001 ****.

To confirm this effect *in vivo*, we immunized mice with P3 or with peptide SEB, designed as in Fig. 6b. We found that P3 induced a broad variety of cytokines in peptide immunized spleen (Fig. 6b-c). The transcription levels of those cytokines (IFN-γ, GZMB, IL6, TNF-α, IL-1β) were increased in spleen of P3-treated mice (Fig. 6b). By staining splenocytes with anti-CD8, we also found that P3 induced in increase in the percentage of CD8+ expressing IFN-γ (22%), GZMB (35%), IL-6 (5%), TNF-α (5%), IL-1β (35%) as compared to control (no peptide immunized group), while SEB immunized group did not show any effect (Fig. 6c).

An increasing frequency of T effector memory (TEM, CD44+CD62L-) was also seen in P3 and SEB peptide-treated mice (Fig. 6c). In addition, we found P3 also induced inflammatory circulating response by increase percentage of CD8+ expressing IFN-γ (20%), GZMB (20%), IL-6 (30%) as compared to control (no peptide immunized group), while SEB immunized group did not show any effect in blood (Supplementary Figure 9). These data suggest that P3 is not only capable of inducing T cell proliferation with an increase in molecules associated with effector function but also can induce inflammatory cytokines in T cells.

### S2 P3-specific CD8+ T-cells TCR profiling

Since we showed that P3 induced responses in 25-30% of T-cells, we also examined the TCR repertoire of the expanded CD8-T-cells. Human PBMCs from 3 individual donors were stimulated with P3 for 6 days, CD8+ T-cells were isolated and 200ng RNA from CD8+ T-cells were sequenced. The raw sequencing data for each sample were mapped to germline segments using *mixcr* (MiLaboratory, version 3.0.11), to generate a clonotype list where a unique combination of V and J segments and the CDR3 nucleotide sequence define each entry. The TCR repertoire NGS results were then analysed with packages *dplyr* and *tidyr* in the environment of R (4.4.1). The co-expressing α and β chain clones by three individuals were counted for clonal fraction and diversity analyse. Top 10 expressing clones for TRAV and TRBV were visualised with *ggplot*. Collectively, we obtained 138538 TRAV and 94767 distinct TRBV clonotypes with P3 stimulation compared with 215209 TRAV and 81943 TRBV clonotypes in CTL samples. We first measured the diversity and clonality of P3 stimulated samples when compared to control samples (n=3). Both the Gini-Simpson and D50 diversity indices were higher TRBV clonotypes in P3 stimulated samples, indicating there were a more diverse TRBV repertoires with P3 stimulation (Fig. 7a and b). Conversely, there is similar the diverse TRAV repertoires between P3-stimulated and control samples (Fig. 7a and b). The Gini coefficient only showed lower TRBV clonotypes in P3-stimulated samples as compared to CTL and the value is close to 0, so it indicated there was equality of TRBV clonotype frequency in P3-treated samples. (Fig. 7c), no difference in TRAV clonotype frequency. Overall, the P3 induced higher diversity and lower clonality than repertoires of no stimulation samples, as expected for a super antigen. From a panel of 630 alpha chains, four gene segments were seen to increase, TRAV13-1*00 (1404, 1410), TRAV17-1*00 (1406) and TRAV20 (1368) (Fig. 7d). Similarly, from a panel of 800 beta chains, only 4 gene segments TRBV11-2*00 (1437) and TRBV7-9*00 (1390, 1440, 1452) were seen to increase (Fig. 7e). These data are consistent with the expected expansion of a limited number of TCR chains as seen with super-antigens.

**Fig. 7.**
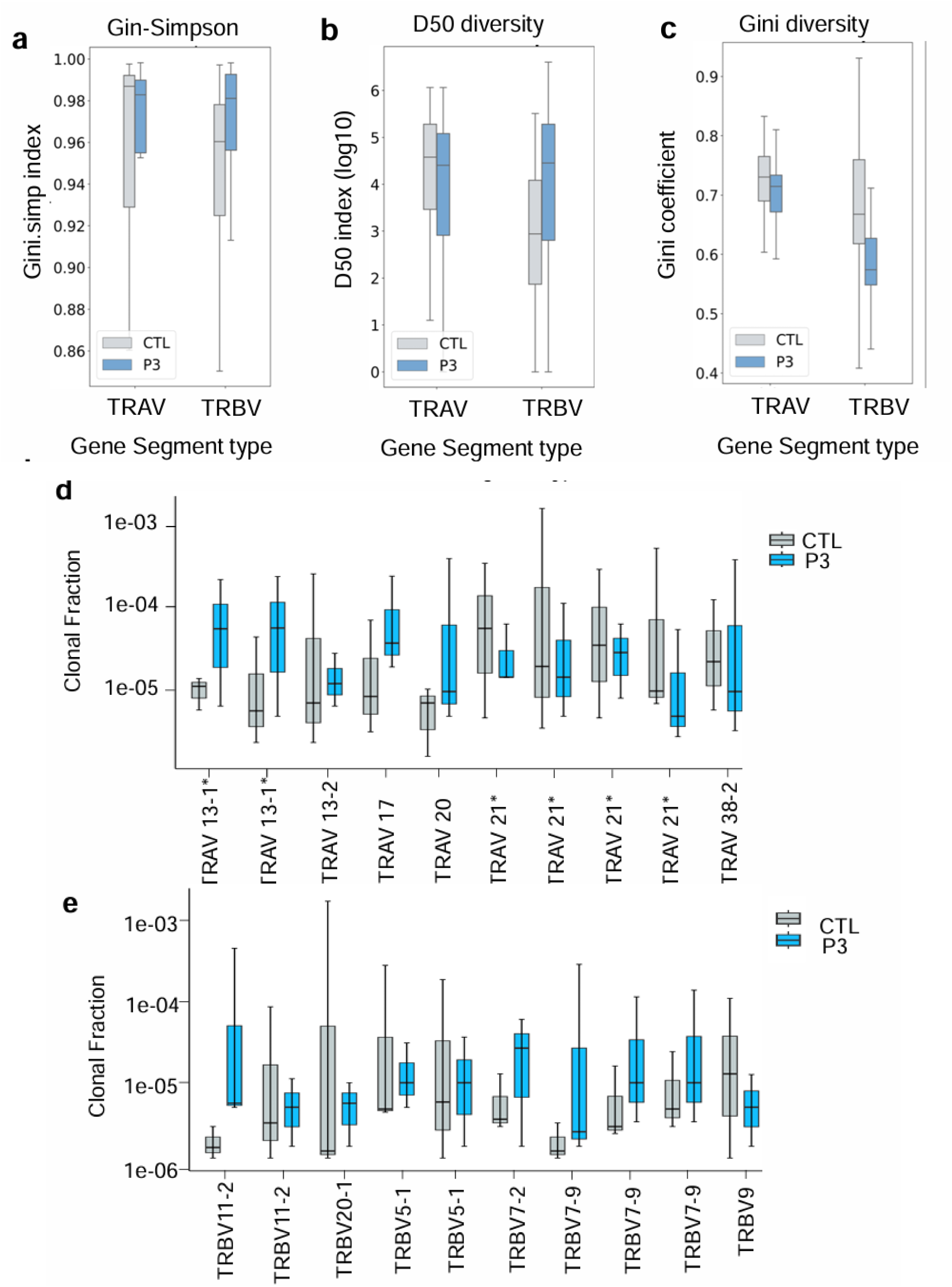
TCR profiling of S2-derived peptide activated CD8+ T-cells. Human PBMCs were isolated from peripheral blood of three healthy donors and were incubated with peptide 3. After 6 days, CD8+ T-cells were isolated and following the TCR repertoire profiling. **a-c** Gini-Simpson (a), D50 diversity (b) and Gini diversity (c) significance analysis were performed to compare the diversity of TCR repertoires between CTL and P3 treated cells. **d-e** The concatenated TCR repertoire NGS results from 3 human PBMCs donors were analysed with packages *dplyr* and *tidyr* in the environment of R (4.4.1). Co-expressing α and β chain clones by three individuals were counted for clonal fraction and diversity analyse. Top 10 expressing clones for TRVA and TRVB were visualised with *ggplot*.c. **d:** Common and distant repertoire results for the TCR-alpha chain. **e:** Common and distant repertoire results for the TCR-beta chain. Data was analyzed and presented as mean ± SEM from 3 healthy human PBMCs donors.

### Peptide similarities to endogenous proteins

These findings identified a small peptide with the capacity to induce unexpected proliferation of T-cells, we next applied the BLAST program to search for homologies of P3 and P3b with the gene databank (Supplementary Table 4). This search revealed extensive homologies with regions within cancer-associated gene 1 protein, E3 ubiquitin ligase TRIM35 isoform 2, MAP3K7 C-terminal-like protein isoforms 2, 3, X1, 7, 6 and 1 as well as prolactin-releasing peptide receptor and others. This raises the potential for the discovery of related endogenous peptides that could stimulate T-cell proliferation associated with autoimmunity and inflammation in the future.

## Discussion

Given the similarity of the cytokine syndrome of TSS and COVID-19, we and others have hypothesized the existence of sequences in the SARs CoV2 virus with homology to SEB/SEE/SEH family. Here, we have identified a sequence (P3) in the Spike 1 (and related P3b) with sequence similarity to various bacterial super-antigens. Virtual docking analysis showed that P3 and P3b bind to MHC and TCR residues in a region that partially overlaps with sites bound by SEB and SEH peptides. Furthermore, we observed that the P3 peptide stimulated a moderate activation of 25-30% of CD4+ and CD8+ T-cells. This stimulation resulted in the upregulation of various activation markers and the increased transcription of the pro-inflammatory cytokine IL-6. Related P3b also induced low level activation of mouse T-cells. The super antigenic nature of P3 was further evidenced by its selective expansion of T-cells expressing specific TCR Vα and Vβ chain repertoires. Lastly, in vivo studies in mice revealed that P3 administration led to elevated levels of proinflammatory cytokines IL-1β, IL-6, and TNF-α. Surprisingly, P3 exhibits homology with naturally occurring mammalian proteins, raising the possibility that peptides derived from these proteins may contribute to autoimmune and inflammatory processes.

Our first observation was that P3 shared approximately 25% homology with super-antigens known to cause TSS ^27,29,30^. Computational docking analysis revealed that P3 binds to overlapping site on MHC class I and class II antigens and the T-cell receptor (TCR) bound by SEB and SEH. For example, P3 and SEH shared three binding residues on TCRα (Ser94, Gln95, Gly52) and two on TCRβ (Arg98, Tyr101). In our initial search, we first identified the unusual DPLQFPELDSFKEEL (P3) variant with the sequence DPLQFPELDSFKEEL; however, later sequences showed the sequence DPLQPELDSFKEEL (termed P3b) which differed by the presence of a single phenylalanine (F) is more frequently found in SARs CoV2. Using CABSDOCK and HPEPDOCK web servers, we found that both peptides bind to a similar region between MHC class II and I and the TCR, although P3b has fewer contact sides consistent with its less effective activation of T-cells.

Consistent with its super-antigen-like properties, P3 and P3b induced the proliferation of 25-35% of CD4+ and CD8+ T-cells, higher than the expected 0.0001-0.001% induced by conventional peptide antigens^31^. Further, a focused TCR repertoire was expanded upon P3 stimulation, with significant expansion of TRAV13-1\**00, TRAV17-1*\*00, TRAV20, TRBV11-2\**00, and TRBV7-9*\*00 gene segments. This selective expansion of specific TCR Vα and Vβ chains is consistent with the super antigenic nature of P3, as previously reported for SEH^52^. P3 stimulated multiple rounds of cellular division (≥5 cycles) in both CD4+ and CD8+ T lymphocytes, accompanied by upregulation of the proliferation marker Ki67. Notably, P3 showed a superior CD8+ T cell stimulatory capacity when compared to the smaller peptides SEB and SEH. Analysis of activation and effector function markers revealed distinct but partially overlapping populations within viSNE and SPADE clustering patterns. Granzyme B expression was confined to a restricted CD8+ T cell subset, displaying more limited distribution than CD69. SPADE analysis identified a responsive CD8+ T cell population characterized by concurrent expression of CD69, IFN-γ, granzyme B, and Ki67, likely representing the primary effector population in the antiviral response. The observed IFN-γ production may be particularly significant given its known paracrine effects on T cell function^53^. In a similar context, the SEB superantigen can activate CD8 T-cells in vivo due to direct TCR engagement and by bystander effects^37^.

P3 upregulated the transcription of pro-inflammatory cytokines IL-6, IL-1β, and TNF-α in vitro and in vivo mouse models. In severe diseases, lymphopenia is correlated with high levels of IL-6 and TNF-α ^9, 10, 48, 49^. Further, high levels of IL-6 are also associated with a poor prognosis in COVID-19, where antibody against IL-6 (tocilizumab) can shorten the hospital stay of COVID-19 patients^50, 51^. A limitation of our study is that despite its ability to induce an inflammatory response, it’s role in COVID-19 remains unclear. It is unlikely to be presented as part of a larger peptide sequence since intact or denatured S1 failed to activate cells in other studies ^54^. Instead, if used, it would be expected to come from the normal processing of the S1 peptide by the proteosome as has been classically demonstrated for other virally derived peptides^55^.

In this context, paradoxically, P3 also acted to rescue the poor response of T-cells taken from patients with a severe disease in terms of an increased expression of CD69 and IFN-γ (Supplemental Figure 10). These results suggest that despite its proinflammatory effects, in certain contexts, the induced CD8+ T-cells responses induced by peptide 3 might be beneficial for development T cell peptide vaccines. Studies have claimed a decrease capacity to produce IFN-γ production in CD8+ T-cells with COVID-19 ^56, 57, 58^.

Sequence homology analysis revealed that P3 shares similarities with various endogenous proteins, including E3 ubiquitin ligase TRIM35 isoform 2 and MAP3K7 C-terminal-like protein. This finding suggests that P3, or related endogenous peptides derived from endogenous proteins, may contribute to the induction of general inflammatory responses in humans. Future studies will investigate the specific cleavage and processing of these peptides by antigen-presenting cells to elucidate their role in immune activation.

## Methods

### Molecular docking of peptide-protein (HLA (I and II)-Peptides-TCR) complexes

Using the two highly regarded web-based application tools CABS-dock ^60^ and HPEPDOCK ^61^, the molecular docking was performed for peptide 3 investigated in the present study. After peptide and protein (Complex immune HLA_TCR) preparation step involving reparation of missing residues, the two different docking approaches that are blind docking (i.e., docking executed without confining any grid box for the target protein) and local docking (i.e., defining a specified grid box center for the respective target protein) was carried out for all peptides against HLA (***(I & II***)-TCR complex. Particularly, two different HLAII-TCR complexes, PDB IDs: 2XN9 ^62^ and 4C56 ^63^ were used, 2XN9 and 4C56 were attached with two crystal structure of staphylococcal enterotoxin H and B, respectively, and regarding the complex HLA I-TCR, we used the PDB IDs 3PWP ^64^. For execution of local docking, the grid box was confined at the interaction interface region of the HLAII-TCR complex or at the region where the enterotoxin B or H attached to the HLAII-TCR complex. The reason behind using two different crystal structure complexes for molecular docking study was raised from the fact that the two the co-crystalized ligand are different (SEH, SEB) and both of them was used in our biological test investigations and also in order to compare the finding and explore the propensity of the peptide 3 to SEH or SEB.

In particular, for executing the docking study, the CABS-dock is a robust computational tool considered the entire protein–peptide system’s flexibility during docking execution in a reasonable time. The CABS-dock program has been widely employed in a variety of applications, including elucidating binding mechanisms of various peptides, peptide dynamics, protein folding evaluation, and structural conformational investigations ^65^. On the other hand, the HPEPDOCK docking protocol used a hierarchical docking approach that includes quick peptide conformational sampling and ensemble docking of generated peptide conformations against the protein. In a comprehensive study that evaluated the efficiency and accuracy of 14 docking programs was revealed that the performance of HPEPDOCK docking program was best among all tools ^66^ to yield the best predictions of peptide-protein complexes. For both the docking protocol execution, mostly the default settings were used. (note that for the input of HPEPDOCK all peptide was converted to their secondary structure using PSIPRED ^67^). The prediction of protein secondary structure is based on position-specific scoring matrices. The reason behind using the two highly regarded servers CABS-dock and HPEPDOCK, besides the two types of Molecular Docking was to compare and confirm accurately the findings. The schema summarizes all the predicted complex docking for both HLA I and HLA II is summarized in supplementary information Supplementary Figure 1 and 2.

### Homology modeling

To examine the relationship of peptide 3 with the various types of spike proteins of SARS-CoV and SARS-CoV2 WT and key variants of concern (Alpha, Beta, Gamma, Delta and Omicron), we conduct a homology modeling. The different strains of Coronaviruses were retrieved from NCBI database while the variants of SARS-Cov2 were collected from Protein Data Bank PDB ^68^, and the references of strain used are summarized in the tables below. The homology was predicted using the online epitope conservancy analysis server ^69^ of Immune Epitope Database (IEDB). Here information references of all strain of Coronaviruses from NCBI, that were used as input for homology modeling: SARS-Cov_2 (WT) (accession no: NC_045512.2), OC43 (accession no: NC_006213.1), NL63 (accession no: NC_005831.2), HKU1 (accession no: NC_006577.2), 229E (accession no: NC_002645.1). And, PDB IDs of the variants of SARS-CoV 2 used for homology modeling: Alpha_(B.1.1.7) (7R1A), Beta_(B.1.351) (7Q6E), Gamma_(P.1) (7V83), Delta_(B.1.617.2) (7SO9), Omicron_(B.1.529) (7WPA).

### Prediction of HLA allele binding for peptide3

Using NetMHCpan - 4.1 (Vanessa Jutz 2017) web server to study the propensity of our peptide 3 to HLA Class I in order to predict binding of peptide 3 to 54 HLA Class II, alleles (5 HLA-DR alleles, 20 HLA-DQ, 9 HLA-DP) NetMHCII - 2.3 (Jensen KK 2018) web server was used for the purpose ^70^. FASTA format was used in input for both peptide and HLA class I and II.

### Software

We used Pymol and Schrödinger for refinement and complex docking presentation and analyses.

### Isolation and culture human PBMCs

Human peripheral blood samples from uninfected healthy adults were obtained from the Hema-Quebec blood bank where donor informed consent was acquired in accordance with Hema-Quebec’s ethical policies, as approved by the CR-HMR Ethical Approval (Le Comité de protection des animaux du CIUSSS de l’Est-de-l’Île-de-Montréal (CPA-CEMTL), F06 CPA-21061 du projet 2017-1346, 2017-JA-001). All applicable ethical guidelines for research involving human participants were followed. Human PBMCs were isolated from blood obtained from Hema Quebec (Quebec) with no prior diagnosis of or recent symptoms consistent with COVID-19 disease. Clinical data was recorded into standardized case report forms. Freshly PBMCs were rested overnight in cell culture media, include RPMI 1640 medium (Corning, USA), 10% heat-inactivated fetal bovine serum (Gibco, USA), penicillin (100U/ml, Hyclone, USA), at 37°C. On the second day, the PBMCs were cultured without or with peptides (P3, SEB, SEH). After 6 days of culture, cells were resuspended in staining buffer (PBS and 2% FBS) and processed the staining for flow cytometry. All animal experiments were conducted with the approval of the Animal Committee at the Research Center of Hospital Rosemont Maisonneuve, Montréal, Canada. Project Number 2021–2525. Mouse splenocytes were cultured after removal of red blood cells with hypotonic buffer (0.15 M NH_4_Cl, 1 mM NaHCO_3_, 0.1 mM EDTA, PH 7.25) in RPMI containing 10% FCS, 5% glutamine, 5% penicillin/streptomycin and 2-mercaptoethanol at 5 × 10^−5^ M. We have complied to all applicable ethical regulations for animal use.

### T cell proliferation assay

Before culturing with peptides, PBMC were prelabeled with 5 mM Tag-it Violet™ Proliferation and Cell Tracking Dye (Biolegend) in PBS with 0.1% FBS for 10 min in a 37°C water bath. Excessive dye was removed by using 10% FBS and further rinsed with cell culture media. Cells were then resuspended in cell culture media at concentration 2 ×10^6^ cell/ml and plated at 200.000 cells per well in a 96-well round-bottom polystyrene plate at a final volume of 100ul. These cells were cultured with 1ug/ml peptide (P3, SEB and SEH) or media (negative control) or full-length staphylococcal enterotoxin B (positive control) for 6 days. At day 6, cells were prepared for flow cytometry staining.

### Flow cytometry

The day of the FACs staining, after fixable viability stain 510 (BD), PBMCs cells were stained with the following antibodies for 30min at 4°C: AF700-conjugated anti-CD3 (clone: UCHT1), BV785-conjugated anti-CD8 (clone: RPA-T8), BUV395-conjugated anti-CD4 (clone: RPA-T4), BV605-conjugated anti-PD-1 (clone: EH12.1), and APC-Cy7-conjugated anti-CD69 (clone: FN50) in FACs buffer during 30min at 4°C in the dark. After, cells were washed and fixed in Fixation/Permeabilization Buffer during 45 min at 4°C in the dark and finally stained with PEcy7-conjugated anti-IFNg (clone: 4S.B3), PE-CF594-conjugated anti-Granzyme B (clone: GB11), FITC-conjugated anti-Ki67 (clone: 11F6). Cells were analyzed using the Fortessa X-20 flow cytometer.

### Flow cytometry and high-dimensional data analysis

Data were analyzed using FlowJo version 10.5.3 (FlowJo). After doublets removal using the forward-scatter area versus forward-scatter height plot, dead cells were excluded by gating on the BV510–negative cells. Next, CD4+ and CD8+ cells were gated within viable CD3+ lymphocytes and analyzed separately for each functional marker (Supplementary Fig.2). viSNE and SPADE analyses were performed on Cytobank (https://cytobank.org). In this case, CD4+ T-cells, and CD8+ T-cells were analyzed separately. viSNE analysis was conducted with equal sampling, using 3000 iterations, a perplexity of 50, and a theta of 0.5. The following markers were included to generate the viSNE maps: CD8, PD-1, CD69, Ki67, IFNg, and GzmB.

### Mice immunization

Female C57BL6 mice aged eight to ten weeks were divided into three groups (n=6 per group) of the following: control (no peptide injection), SEB peptide injection, and P3 peptide injection. Mice were sensitized by means of intraperitoneal injection of 5ug of the peptide mixed with Aluminium hydroxide (Sigma Chemical, St Louis, Mo, USA) (the peptide: adjuvant ratio is 30:70 v/v) in total volume of 100ul on day 0, 7, 14 and 21. The mice were sacrificed one week after the last administration. The blood and spleen were collected. Blood cells and splenocytes were extracted and stained with surface antibodies, such as AF700-conjugated anti-CD3 (clone: 500A2), BV650-conjugated anti-CD8 clone: 53-6.7, PerCp5.5-conjugated anti-CD4 (clone; RM4-4), BV605-conjugated anti-CD44 (clone: IM7), APC Cy7-conjugated anti-CD62L (clone: MEL-14); and cytokines antibodies, such as anti-PE-eFluor610-conjugated anti-IFNg (clone: XMG1.2), PE-conjugated anti-Granzyme B (clone: NGZB), PECy7-conjugated anti-IL1b (clone: NJTEN3), BV786-conjugated anti-TNFa (clone: MP6-XT22), AF488-conjugated anti-Ki67 (clone: 11F6).

### Quantitative real-time PCR

Total RNA was extracted from the PBMCs and reverse-transcribed into cDNA using All-In-One 5X RT MasterMix (Applied Biological Materials Inc, BC, Canada). Real-time PCR amplification of the cDNA was analyzed using Bright Green 2X qPCR Master Mix-Low ROX (Applied Biological Materials Inc, BC, Canada) and QS12K Flex system (Thermo Fisher Scientific). The results were analyzed using CFX Manager software and normalized to the expression levels of the housekeeping gene 18S. The primers for qPCR analysis are shown, human-18S-F (GATTAAGTCCCTGCCCTTTGT), human-18S-R (GTCAAGTTCGACCGTCTTCTC), human-IL1b -F(GAAGCTGATGGCCCTAAACAG), human-IL1b -R (AGCATCTTCCTCAGCTTGTCC), human-IL6-F (AGACAGCCACTCACCTCTTCA), human-IL6-R (CACCAGGCAAGTCTCCTCATT), human TNFa -F (GTGCTTGTTCCTCAGCCTCTT) and human TNFa -R (ATGGGCTACAGGCTTGTCATC).

Total RNA from spleen from peptide immunization mice were obtained and run RT-PCR, same as previously described. The results were analyzed using the CFX Manager software and normalized to the levels of the housekeeping gene b-actin. The primers for qPCR analysis are shown, mouse-b-actin-F (TCTTTGATGTCACGCACG), mouse-bactin-R(TACAGCTTCACCACCACA), mouse-IFNg-F (ATCTGGAGGAACTGGCAAAA), mouse-IFNg-R(TTCAAGACTTCAAAGAGTCTGAGGTA), mouse-TNFa -F(TGGCCTCCCTCTCATCAGTT), mouse-TNFa-R(TCCTCCACTTGGTGGTTTGC), mouse-GzmB-F(CCATCGTCCCTAGAGCTGAG), mouse-GzmB-R(TTGTGGAGAGGGCAAACTTC), mouse-IL6-F(GGGACTGATGCTGGTGACAA), mouse-IL6-R(ACAGGTCTGTTGGGAGTGGT), mouse-IL1b -F(CTGGTACATCAGCACCTCFCA), mouse-IL1b -R(GAGCTCCTTAACATGCCCTG).

### TCR sequencing

PBMCs were cultured with P3 peptide for 6 days. CD8+ T-cells were extracted and processed for bulk TCR sequencing. TCR repertoire profiling was conducted using the SMARTer TCR α/β Profiling Kit (Takara Bio, USA), following the manufacturer’s protocol. RNA was isolated using the Qiagen RNA isolation kit. 200ng RNA from antigen induced or unstimulated PBMCs was subjected for preparing TCR cDNA libraries by using the SMARTer Human TCR a/b Profiling Kit (Takara Bio USA, Inc). The kit utilizes SMART technology (Switching Mechanism At 5’end of RNA Template) combined with 5’RACE to capture the entire V(D)J variable regions of TCR transcripts, followed by two rounds of semi-nested PCR to generate TCR-α and the β-chain libraries. The libraries were then sequenced using Illumina next-generation sequencing (NGS) platform. Sequencing was carried out using a 600-cycle MiSeq Reagent Kit v3 (Illumina, Inc.) with 2 x 300 base pair reads. The NGS data were analyzed using MiXCR online tool.

### TCR repertoire analysis

TCR repertoire analysis was performed using Python program. The diversity of TCR repertoires was evaluated by the Gini, Gini-Simpson and D50 diversity coefficients. The Gini coefficient measures the inequality in the frequency distribution of clonotypes, which values close to zero expressing full equality of clonotypes frequencies, while a Gini coefficient of 1 reflects maximum inequality between clonotype frequencies. The D50 coefficient calculates the minimum number of distinct clonotypes that account for at least 50 percent of sequencing reads obtained after amplification and sequencing. The Gini-Simpson index presents the probability of interspecific encounter. The graphs of the TCR alpha chain and beta chain were drawn with packages *dplyr* and *tidyr* in the environment of R (4.4.1), Co-expressing α and β chain clones by three individuals were counted for clonal fraction and diversity analysis. The top 10 expressing clones for TRVA and TRVB were visualized with *ggplot*.

### Ex vivo culturing of PBMCs from COVID-19 patients

In line with the McGill University Health Centre Research Institute (MUHC-RI) Ethics Board approval of the human study (#2021-6081), informed consent was obtained from patients admitted to the McGill University Health Centre (MUHC) with PCR confirmed SARS-CoV-2 infection between April 2020 and March 2021 (n=5). Peripheral blood samples (K2EDTA-preserved whole blood and/or sera were obtained from this patient cohort and healthy adult donors (HDs) with no prior diagnosis of or recent symptoms consistent with COVID-19 disease. Clinical data were recorded into standardized case report forms. The median time from patient admission to an initial collection of PBMC samples was 3 days (interquartile 1-8 days). Serial samples from a subset of patients were collected at time points out to 36 days from admission. Clinical outcomes and severity of COVID-19 manifestations were assessed using the World Health Organization (WHO)’s COVID ordinal scale. Scores of <=4 were considered to be mild manifestations of disease and >=5 were considered to be severe. Clinical laboratory data were abstracted from the time points closest to that of research blood collection as well as from time points associated with extreme values.

### Statistics and Reproducibility

Statistical analyses were performed using GraphPad Prism software version 7.0 for Windows (GraphPad Software). Results are presented as mean ± SEM of a representative experiment. The sample sizes (n= 4-5) in each figure are indicated in the respective figure legend. Statistical significance differences between peptide-stimulated cells and negative controls were assessed using an unpaired student’s T-test. The significant difference in mean values between the two groups was defined as p ≤ 0.05 *, p ≤ 0.01 **, p ≤ 0.005 *** and p ≤ 0.001 ***.

## Supplemental information

Supplemental tables and figures can be found below.

## Reporting summary

Further information on research design and raw data is available in supplementary files linked to this article.

## Data availability

All data and materials used in the analysis will be available to any researcher for the purpose of reproducing or extending the analyses. The underlying source data for this study can be found in the Supplementary Data file. The RNA-sequencing reads for TCR profiling that support the findings of this study have been deposited in NCBI Gene Expression Omnibus (GEO) under the accession code GSE281135, which includes raw data and processed files including V and J reads. All other data are available from the corresponding author.

## Code availability

The codes that support relevant TCR clonality are provided in GitHub (via Zenodo), at https://doi.org/10.5281/zenodo.14036310.

## Acknowledgments

C.E.R. was supported by the Canadian Institutes of Health Foundation Grant (159912) and National Institutes of Health US grant RO1 AI049466. We thank Drs Nathalie Labrecque, Benjamin Gordon for helpful comments on manuscript.

## Author contribution

C.E.R. designed peptides. T.H.T. conducted most of the experiments; B.F.E. conducted the peptide docking binding site and modeling analysis; N.M. and T.H.T. prepared NGS samples; C.L., A.N. and T.H.T. conducted analysis of the TCR spectrum; N.R., A.G., R.K. and C.M. coordinated access to Covid-19 patients from McGill clinic; C.E.R and T.H.T. and B.F.E. drafted the manuscript; C.M., R.K., D.R. and C.E.R reviewed the manuscript.

