## Supplementary files for "The identification of a SARs-CoV2 S2 protein-derived peptide with super-antigen-like stimulatory properties on T-cells"

**TABLES, FIGURES AND LEGENDS**

**Key resources table. The list of peptides was used in this study.**

The sequences of S2 protein-derived peptide (peptide 3 and peptide 3b) and superantigen-derived peptides (SEB, SEH). Their length and structural organization are summarised in table below.

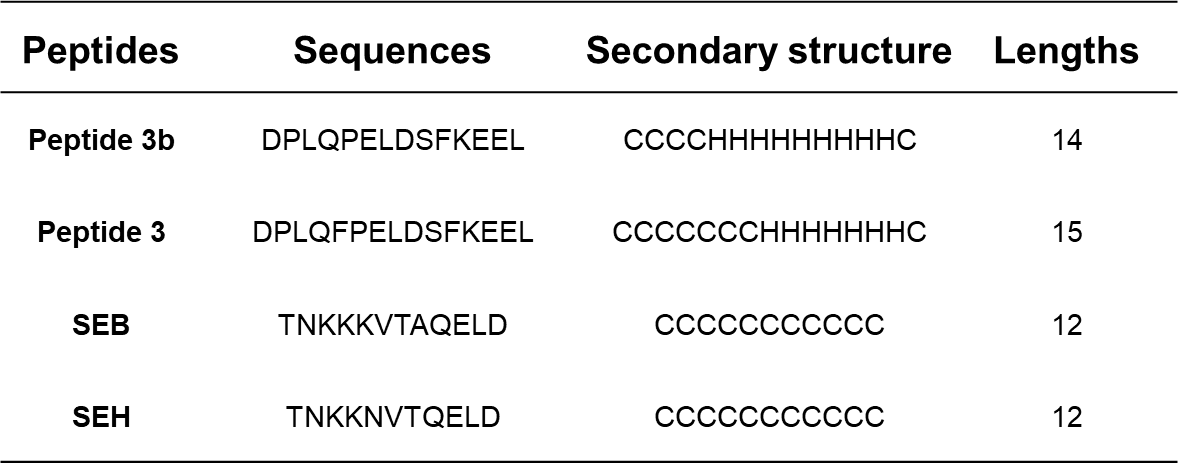

**Supplementary Table 1. The list of list peptide was used in study.**

Homology percentage of S2 protein peptide (peptide 3) to other superantigen (SEA, SEB, SEC, SEE, SEG, SEH, SEI, SEK, SEN).

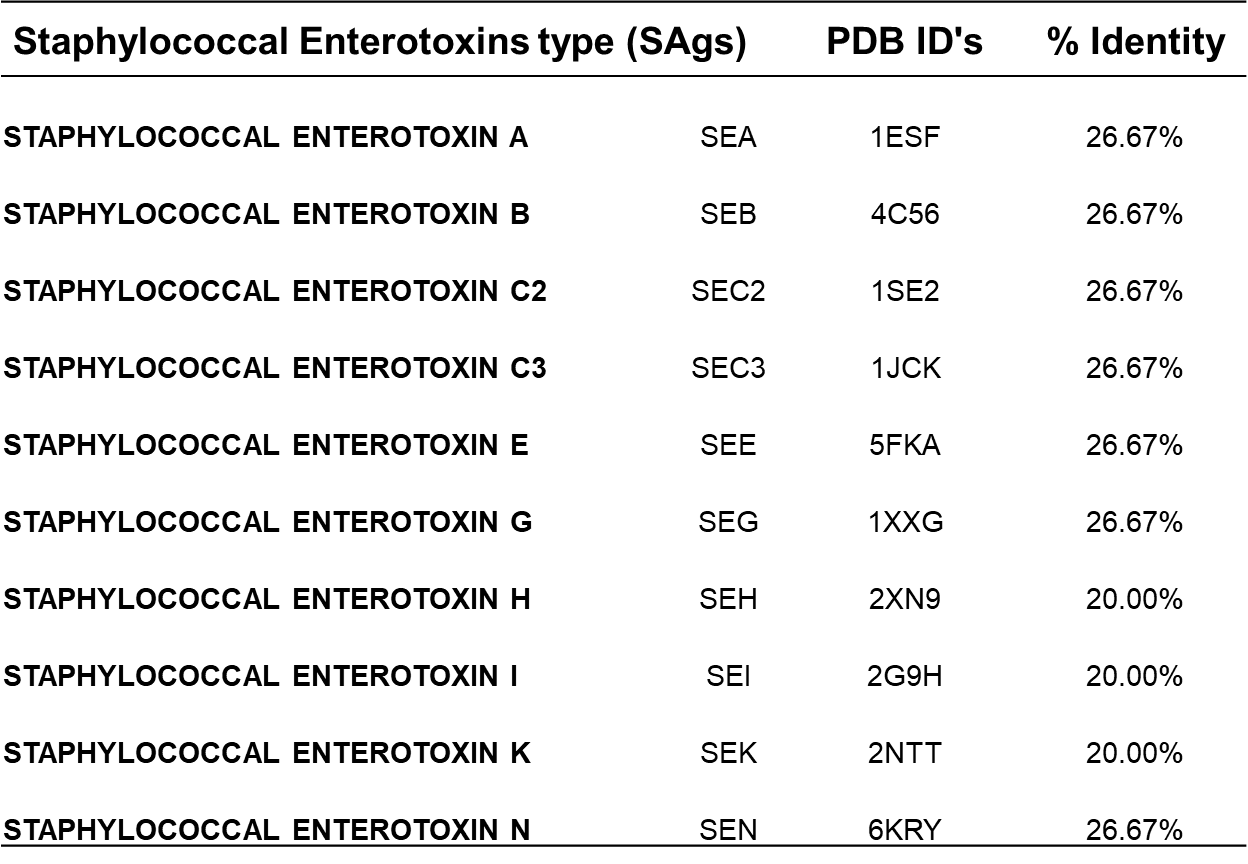

**Supplementary Table 2**. Homology percentage of S2 protein peptide (peptide 3) to other coronaviruses (SARS-CoV, OC43, NL63, HKU1, 229E).

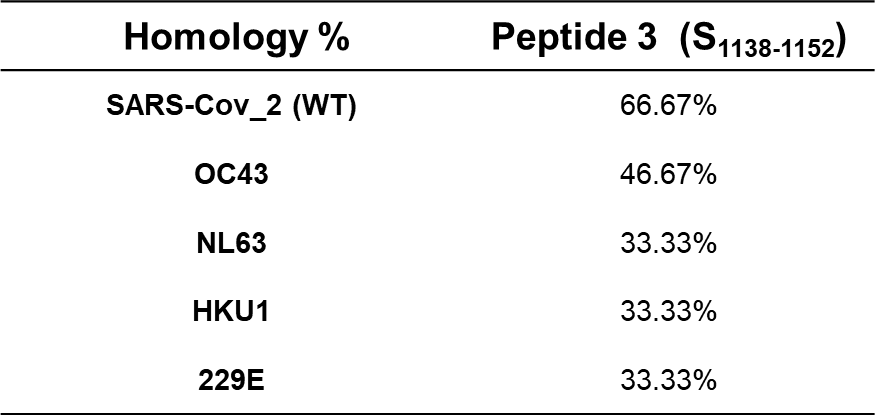

**Supplementary Table 3**. Homology percentage of S2 protein peptide (peptide 3) to other SARS-CoV-2 variants (α, β, γ, δ, and Omicron).

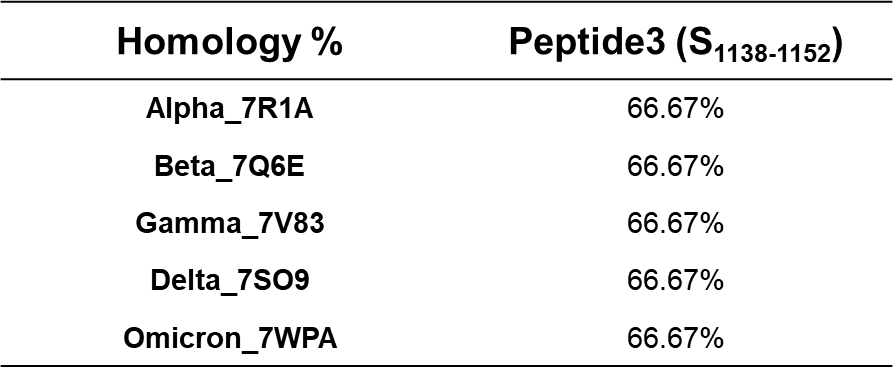

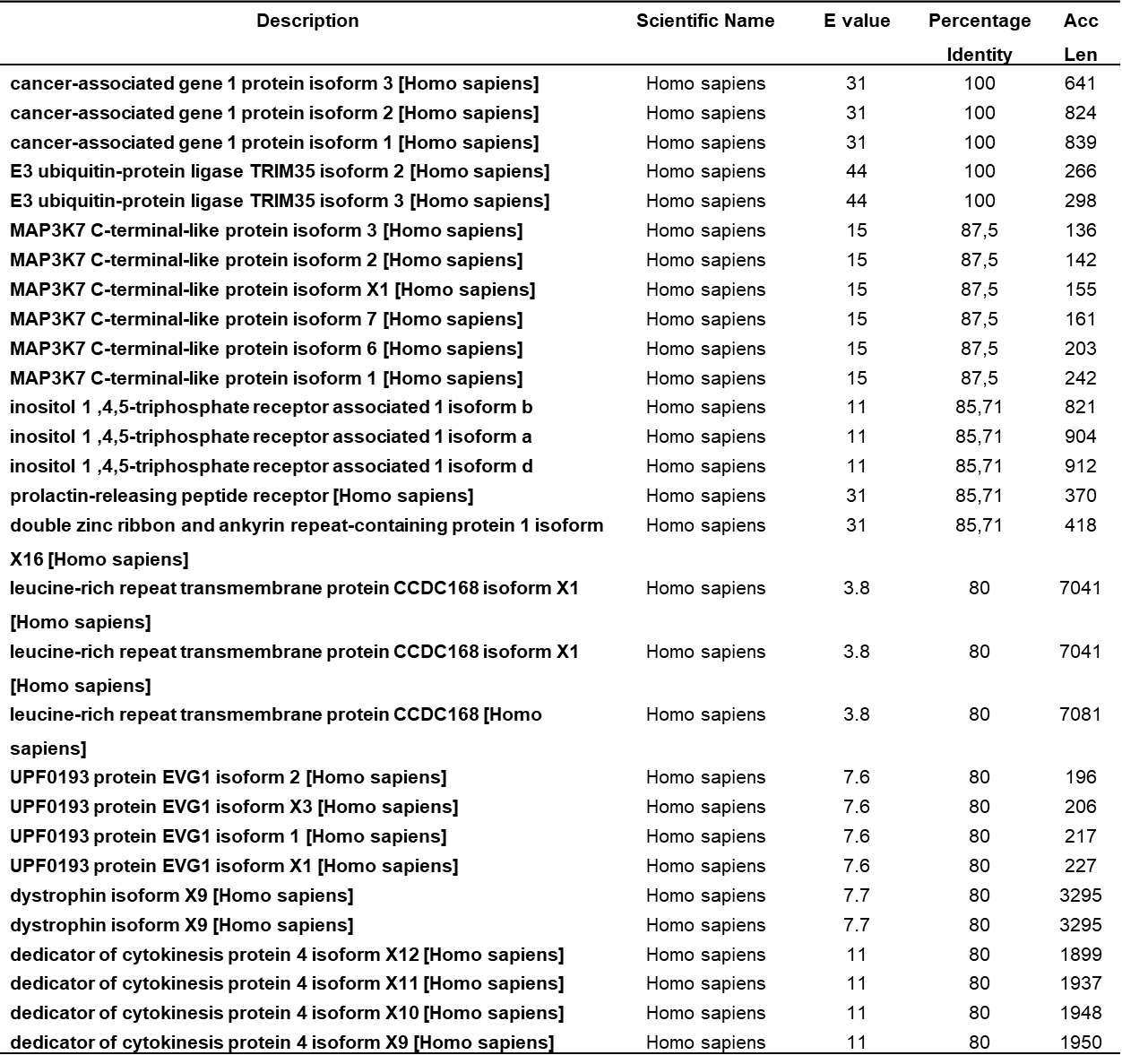
**Supplementary Table 4**. Peptide P3 and P3b share homologous sequences with mammalian proteins.

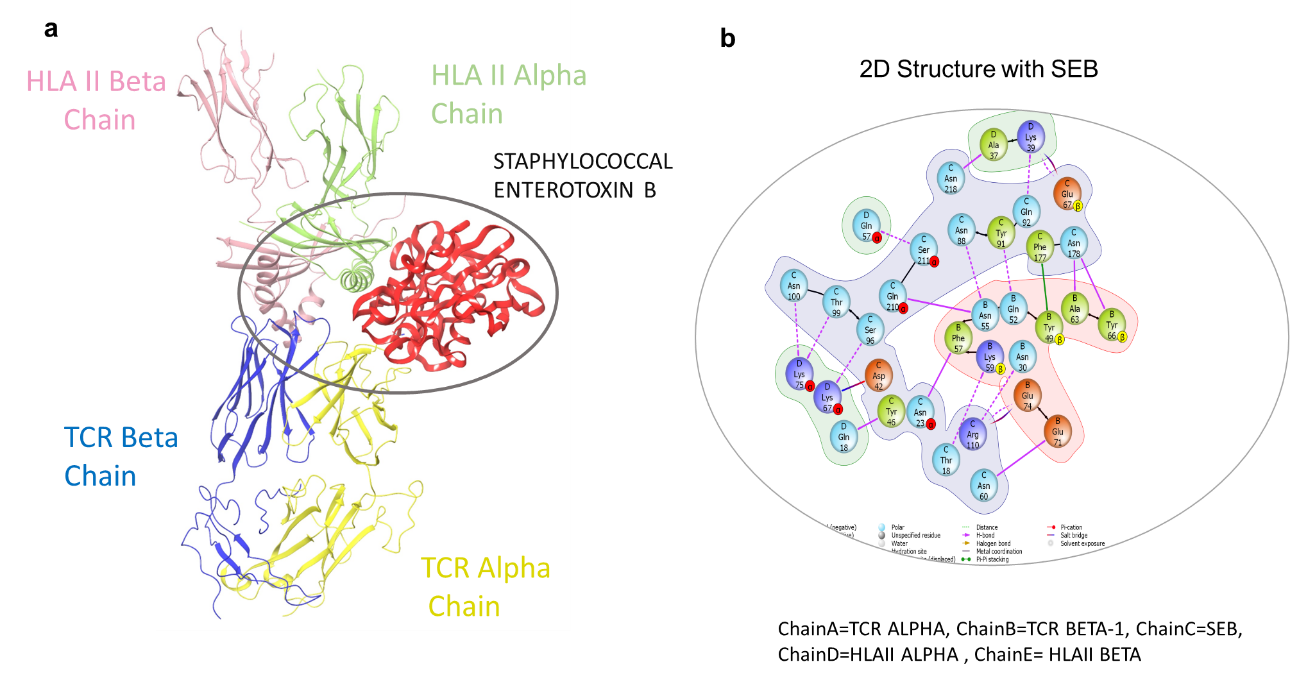

**Supplementary Fig. 1. Overview of three-and two-dimensional structure of the complex docking TCR-HLA-II peptides complex. a** Complex docking of superantigen SEB with TCR and human class MHC (HLA) II antigen displayed in cartoon style for HLA-II_TCR and sAgs SEB is red ribbon, showing binding at the interface connecting TCR to HLA-II. **b** 2D diagram of interaction of sAgs SEB with MHC-class II and TCR residues, the colored diagram shows the binding residues between SEB and TCR and HLA-II.

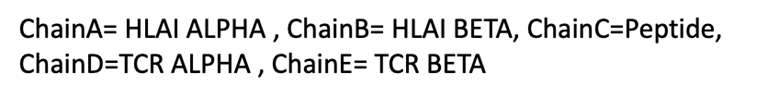

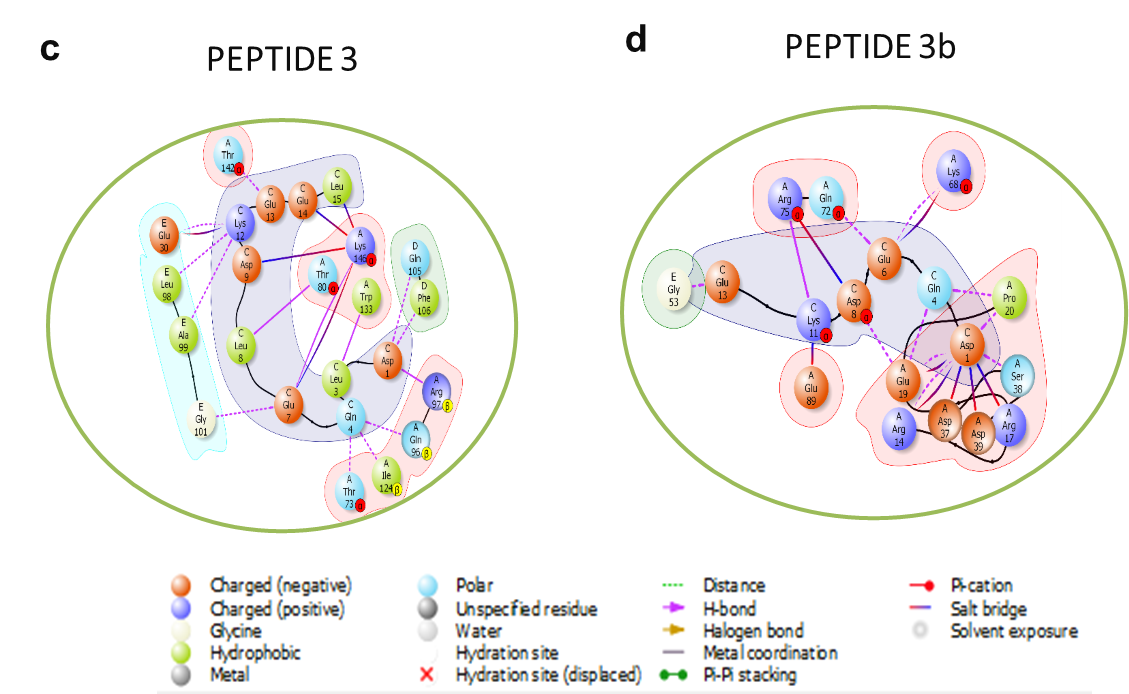

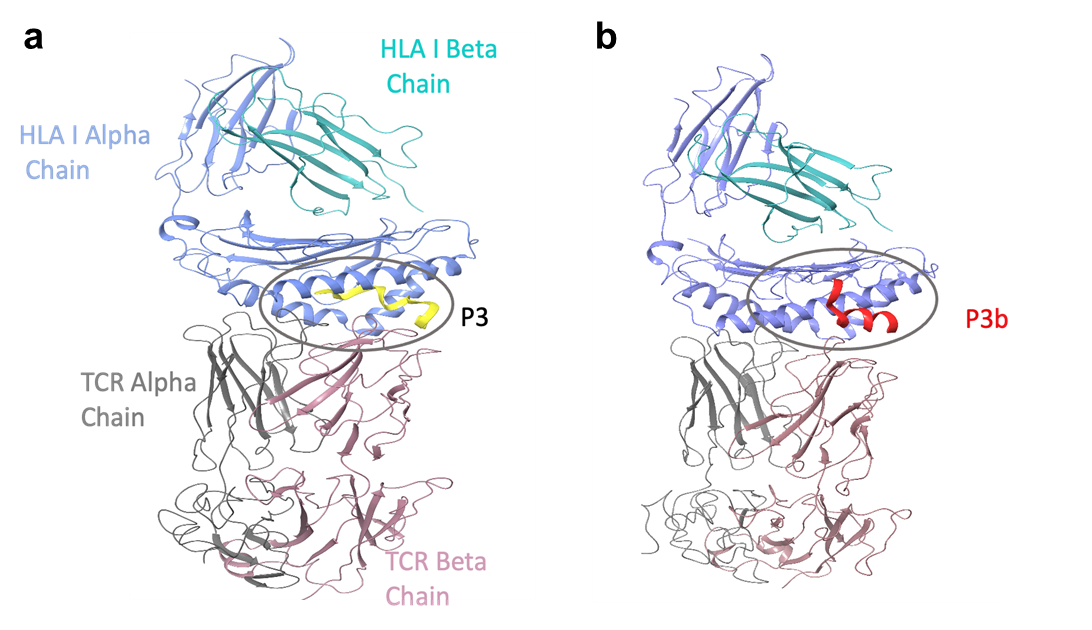

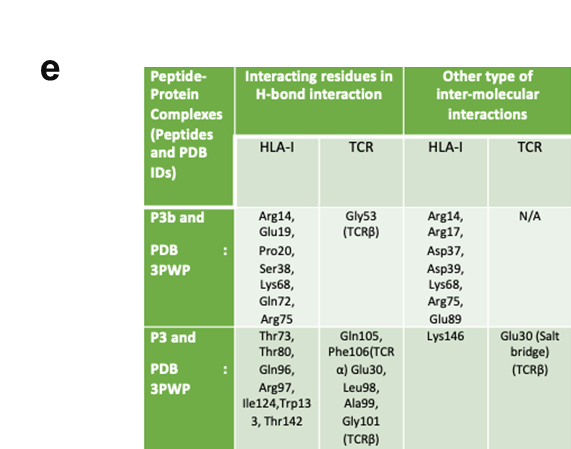
**Supplementary Fig. 2. Overview of three-and two-dimensional structure of the complex docking TCR-HLA-I peptides complex with peptide P3 and P3b. a** Complex docking of P3 with TCR and human class MHC (HLA) I antigen displayed in cartoon style for HLA-I_TCR and peptide P3 is yellow ribbon, showing binding at the interface connecting TCR to HLA-II. **b** Complex docking of P3d with TCR and human class MHC (HLA) II antigen displayed in cartoon style for HLA-I_TCR and peptide P3b is red ribbon, showing binding at the interface connecting TCR to HLA-I. **c** Two-Dimensional interaction view for the best fit conformation of Peptide3 with HLAI-TCR (3PWP) complex showing all type of interactions. **d** Two-Dimensional interaction view for the best fit conformation of Peptide 3b with HLAI-TCR (3PWP) complex showing all type of interactions. **e** Table showing the interactions types(H_bond and other inter-molecular interactions ) residues of both Peptide 3 and Peptide 3b with the complex 3PWP .

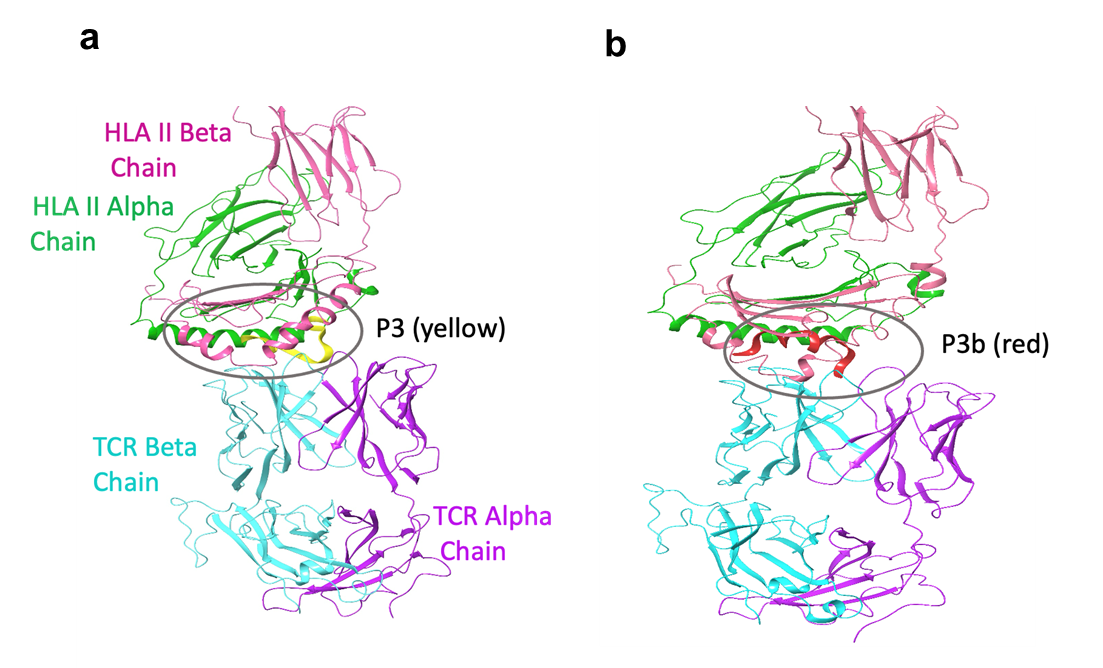

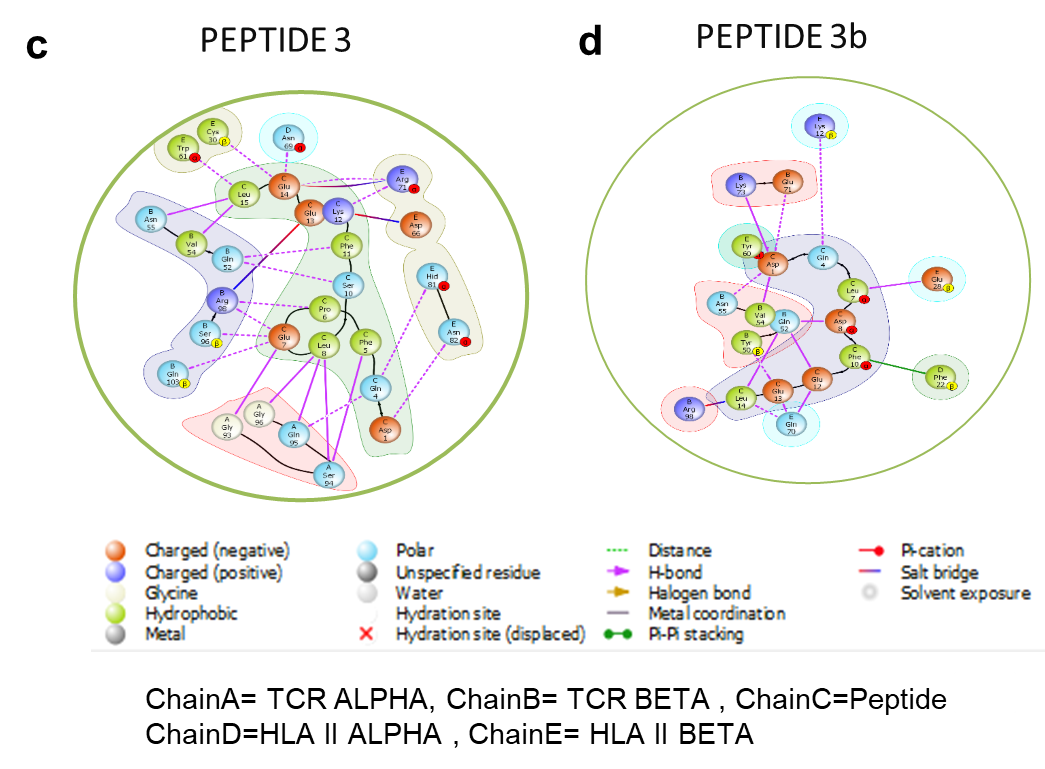

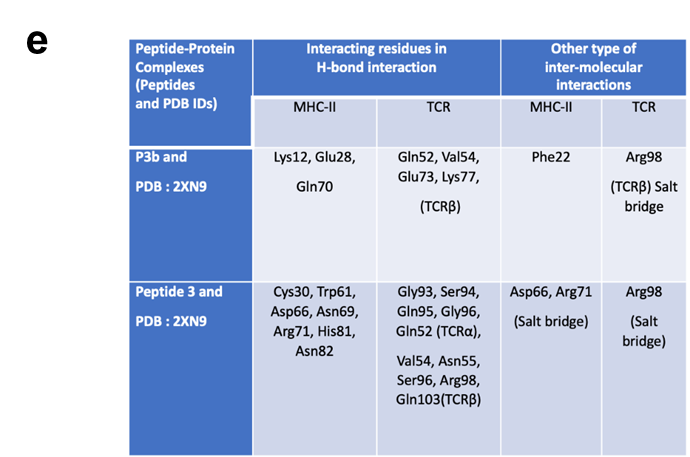

**Supplementary Fig. 3. Overview of three-and two-dimensional structure of the complex docking TCR-HLA-II peptides complex with peptide P3 and P3b. a** Complex docking of P3 with TCR and human class MHC (HLA) II antigen displayed in cartoon style for HLA-II_TCR and peptide P3 is red ribbon, showing binding at the interface connecting TCR to HLA-II. **b** Complex docking of P3d with TCR and human class MHC (HLA) II antigen displayed in cartoon style for HLA-II_TCR and peptide P3b is red ribbon, showing binding at the interface connecting TCR to HLA-II. **c** Two-dimensional interaction view for the best fit conformation of Peptide 3 with HLAII-TCR (2XN9) complex showing all type of interactions. **d** Two-dimensional interaction view for the best fit conformation of Peptide 3b with HLAII-TCR (2XN9) complex showing all type of interactions. **e** Table showing the shared interactions types (H bond and other inter-molecular interactions) residues between both Peptide 3 and Peptide 3b with the complex 2XN9 .

.

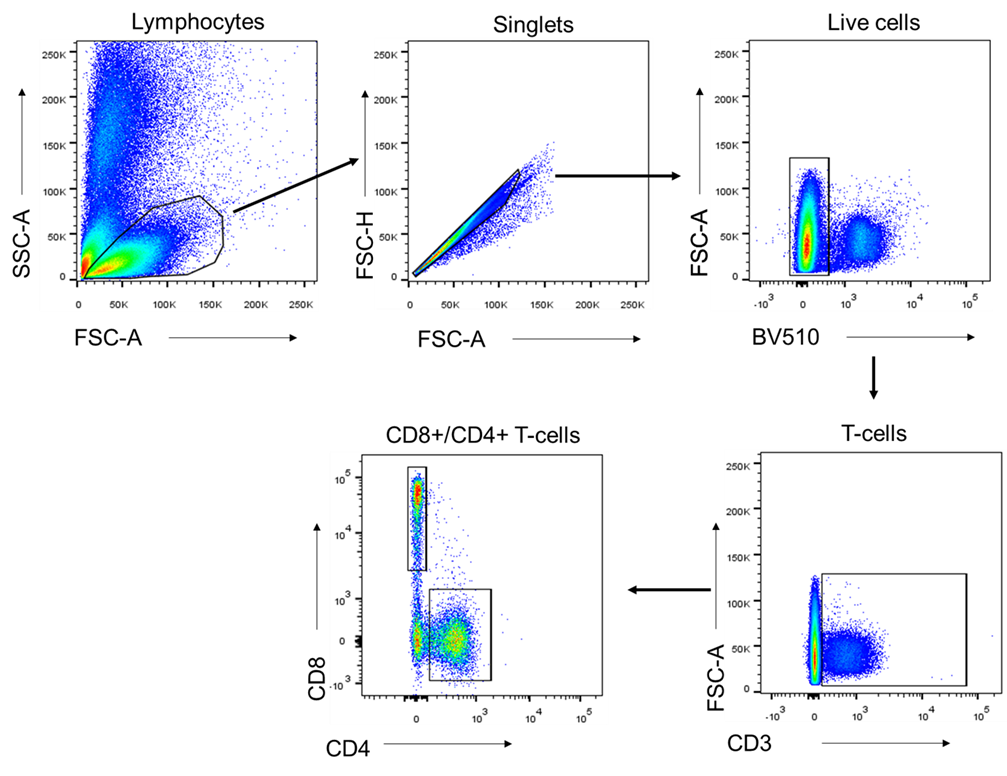

**Supplementary Fig. 4 The gating strategy to phenotype human PBMCs were treated with peptides.**

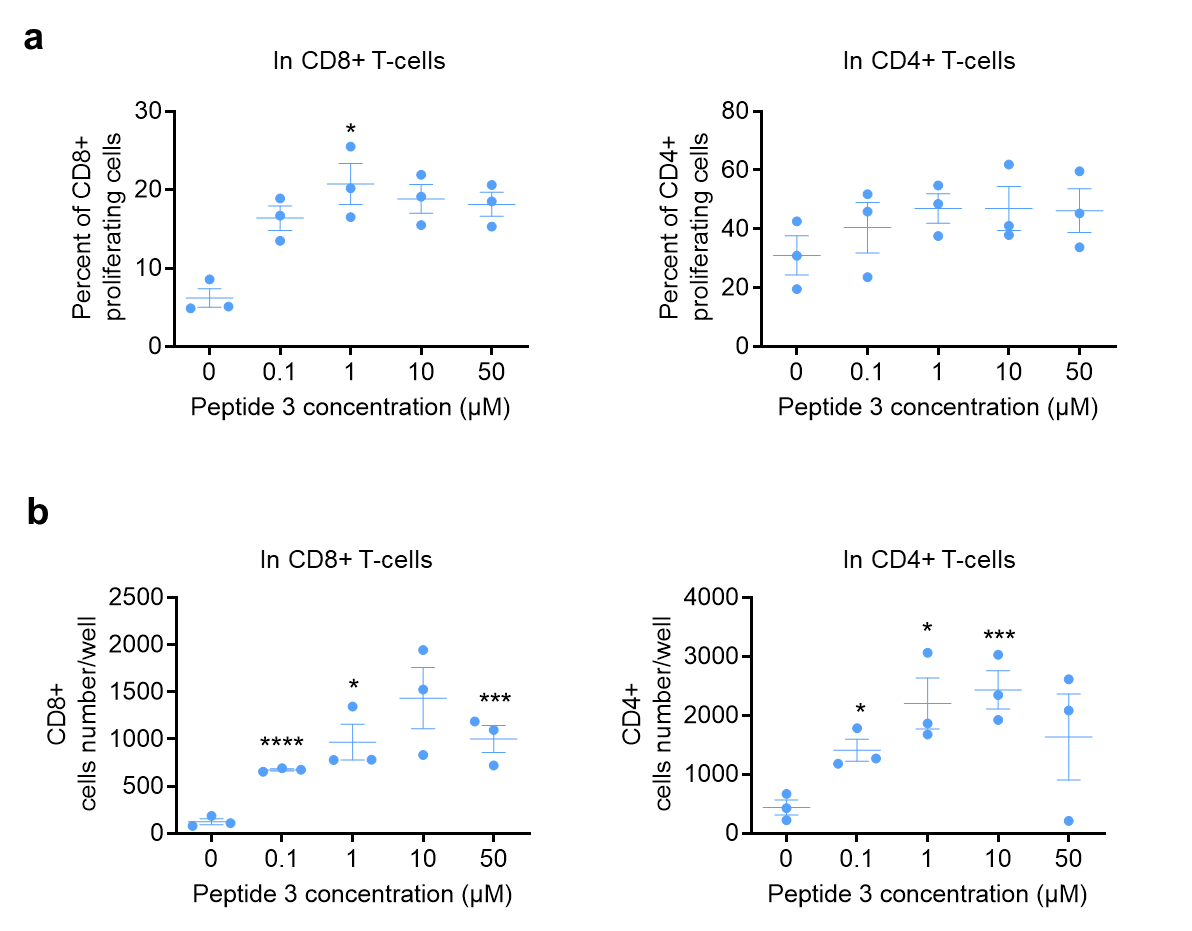

**Supplementary Fig. 5. Proliferative T-cell responses to S2 protein peptide (P3).** PBMCs from health donors were cultured with different concentrations of peptide 3 (0.1, 1, 10, 50 ug/ml) for 6 days. PBMCs cells were stained with Tag-it before incubating with or without peptide 3. The percentage of proliferating cells were measured with Tag-it proliferation assay after 6 days incubation. **a** Percentage of proliferating CD8+ and CD4+ T-cell with different peptide 3 concentrations. **b** Number of proliferating CD8+ and CD4+ T-cell with different peptide 3 concentrations. Data are presented as Mean ± SEM from 3 human PBMCs samples, significant as p ≤ 0.05 *, p ≤ 0.005 *** and p ≤ 0.001 ****.

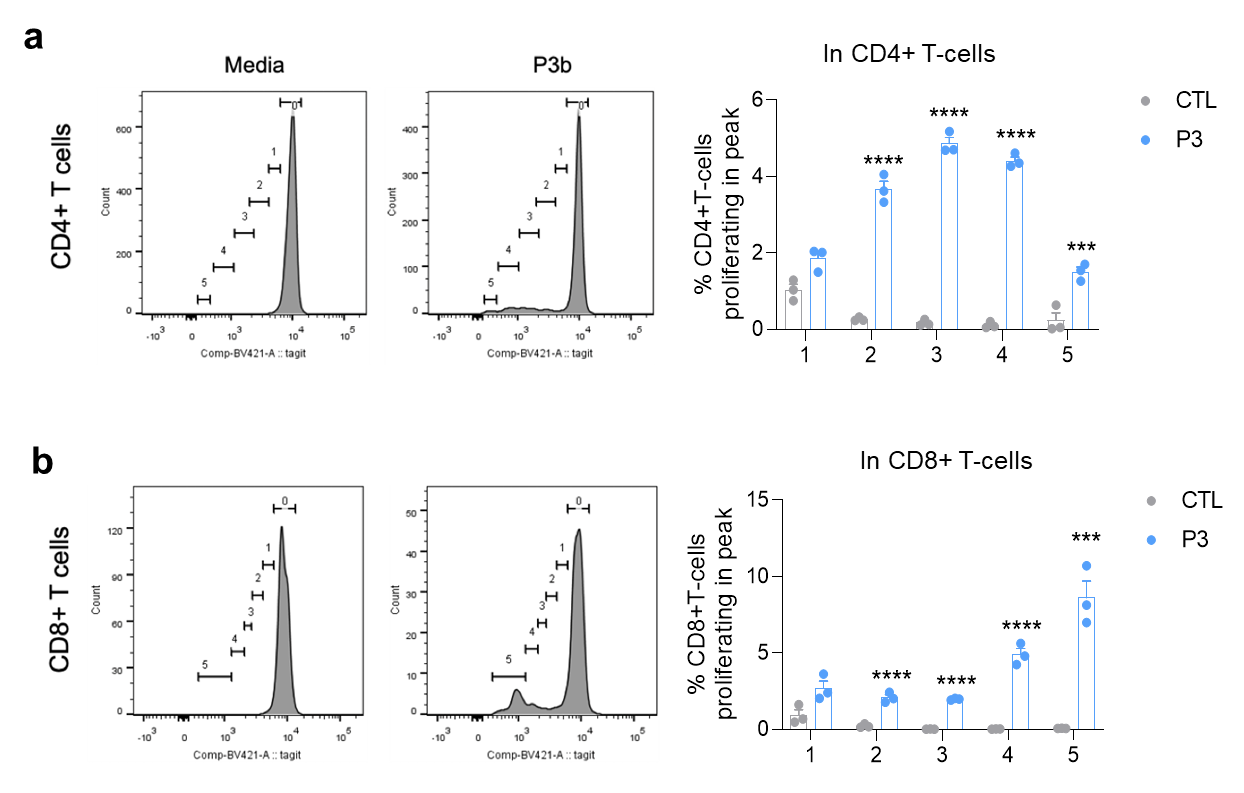

**Supplementary Fig. 6. Proliferative T-cell responses to S2 protein peptide (P3b).** Mouse splenocytes at 2 x 10^6^/ml prelabelled with Tag-it were cultured in RPMI-1640 with 10% FCS and 1ug/ml P3b peptide for 36 hours followed by an analysis of CD4 and CD8 populations by flow cytometry. **a-b** Left panel: Representative flow cytometry plots displaying gating for proliferating cells. The number of divisions was determined in proliferating CD4+ T-cells (a) and CD8+ T-cells (b). Right panel: Histogram showing the percentage of proliferating cells in 5 divisions; CD4+ T-cells and CD8+ T-cells. Data presented as Mean ± SEM from 3 mice, significant at p ≤ 0.005 *** and p ≤ 0.001 ****.

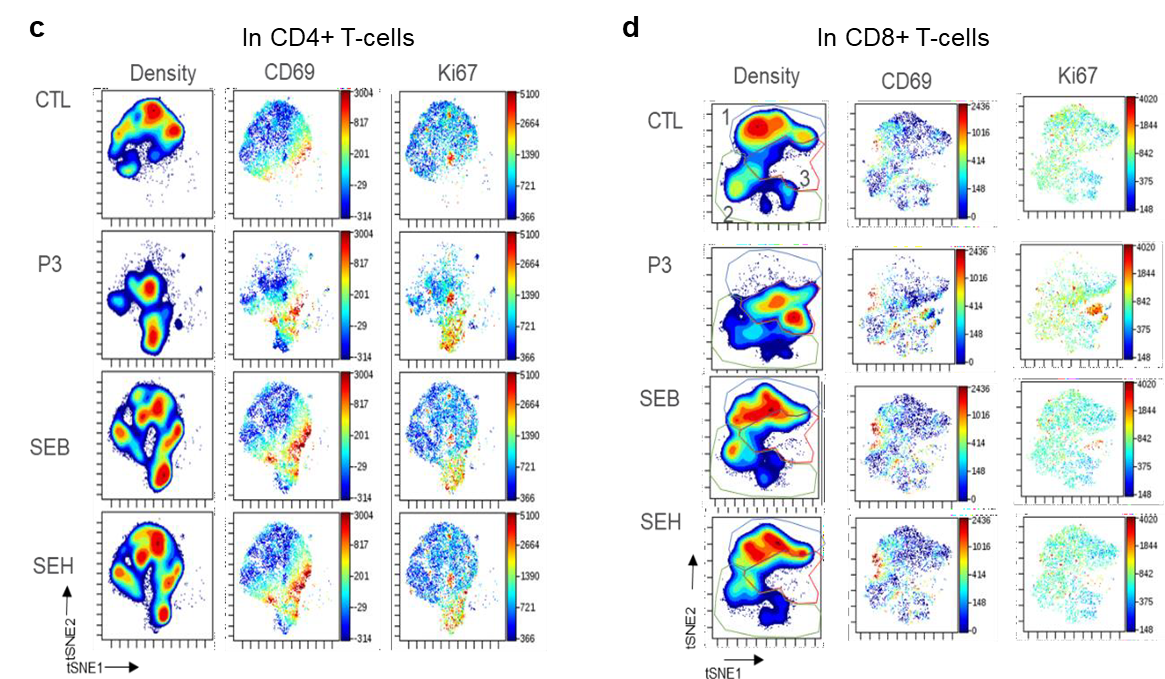

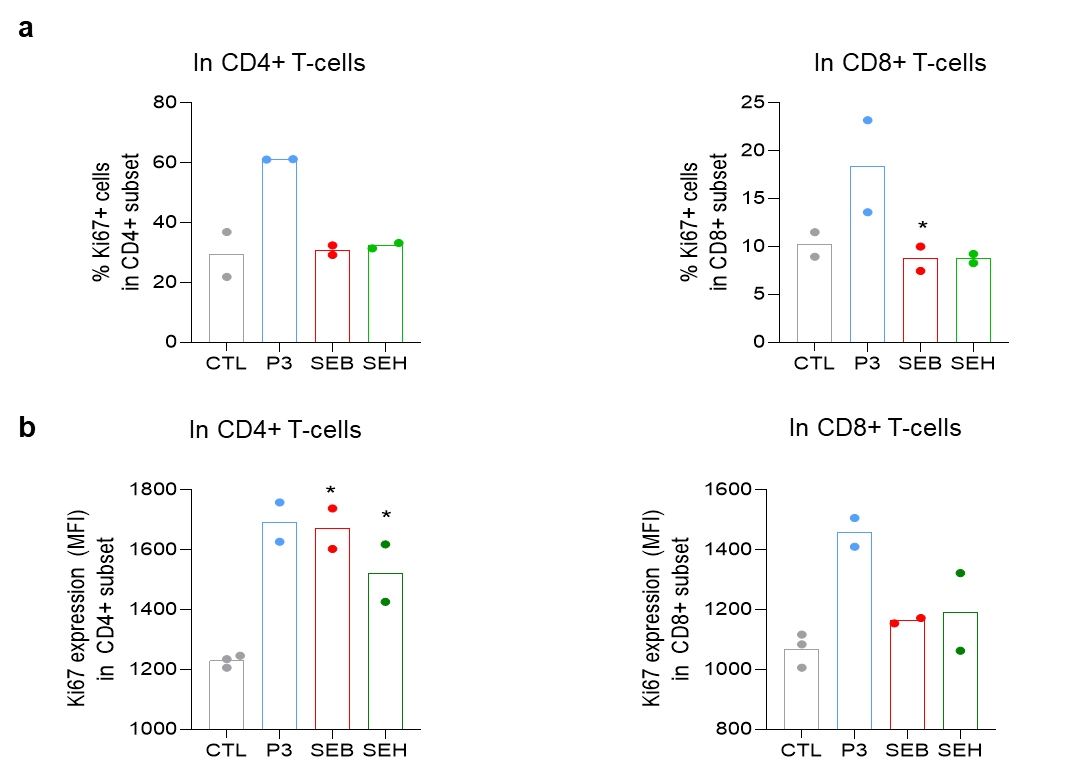

**Supplementary Fig. 7. P3 spike peptide activates CD4+ and CD8+ human T-cells.** Human PBMCs were incubated with peptide 3, SEB and SEH for 6 days, followed by an analysis of activation markers by flow cytometry. **a-b** Ki67 expression on CD4+ T-cells and CD8+ T-cells. Histogram showing the percentage of cells expressing Ki67 (a) and Ki67 MFI expression levels in CD4+ and CD8+ T-cells (b). Data are presented as Mean ± SEM from two human PBMCs samples, significant at p ≤ 0.05 *, p ≤ 0.005 *** . **c** viSNE profiles of CD69 and Ki67 expression on CD4+ T- cells activated by P3 peptide. **d** viSNE profiles of CD69 and Ki67 expression on CD8+ T-cells activated by P3 peptide. The viSNE analysis was run with concatenated samples from 2 human PBMCs samples.

\

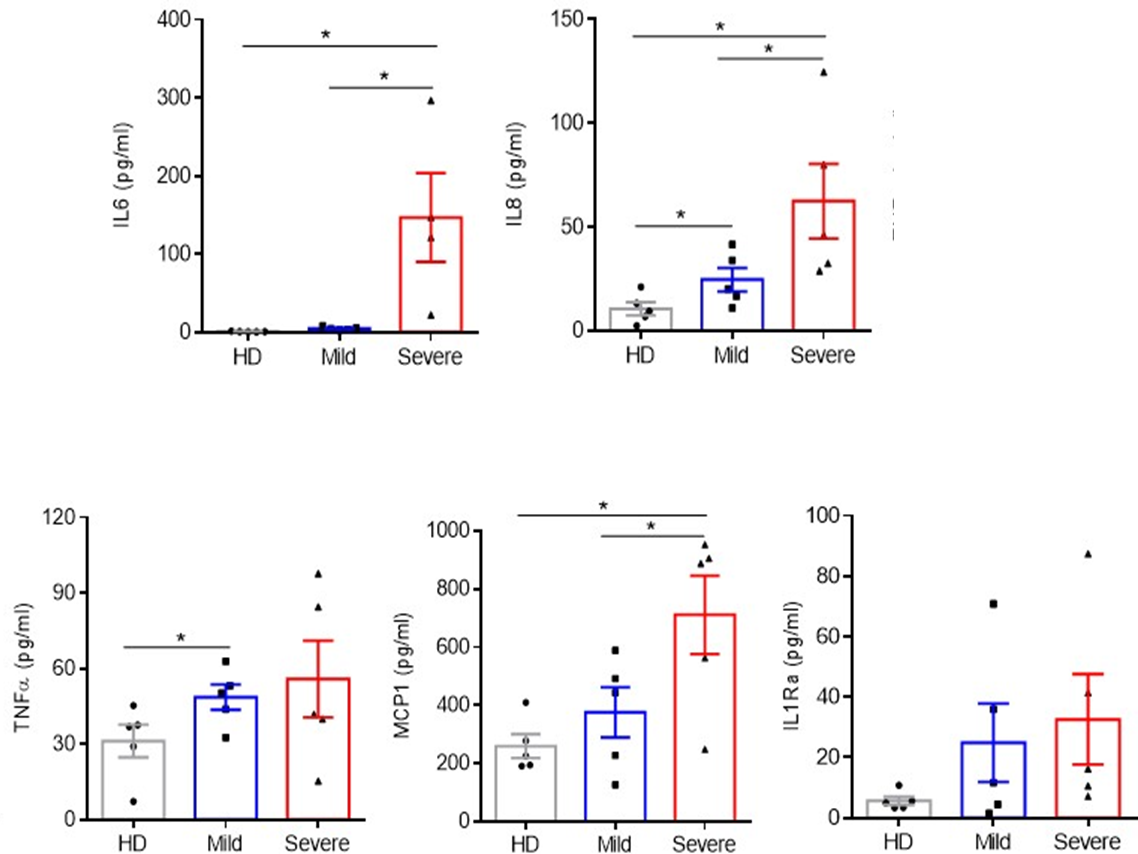

**Supplementary Fig. 8. Profiling serum cytokines in COVID-19 patients’ serum.** Serum concentration of IL6, IL8, TNFa, MCP1 and IL1Ra from heathy donors (n=5) and COVID-19 patients (Mild = 5, Severe = 5). A broad range of cytokines was assessed by using human cytokine array proinflammatory focus 15-plex (HDF15) (Eve Technology, Calgary, AB).The statistic significant at p ≤ 0.05 *.

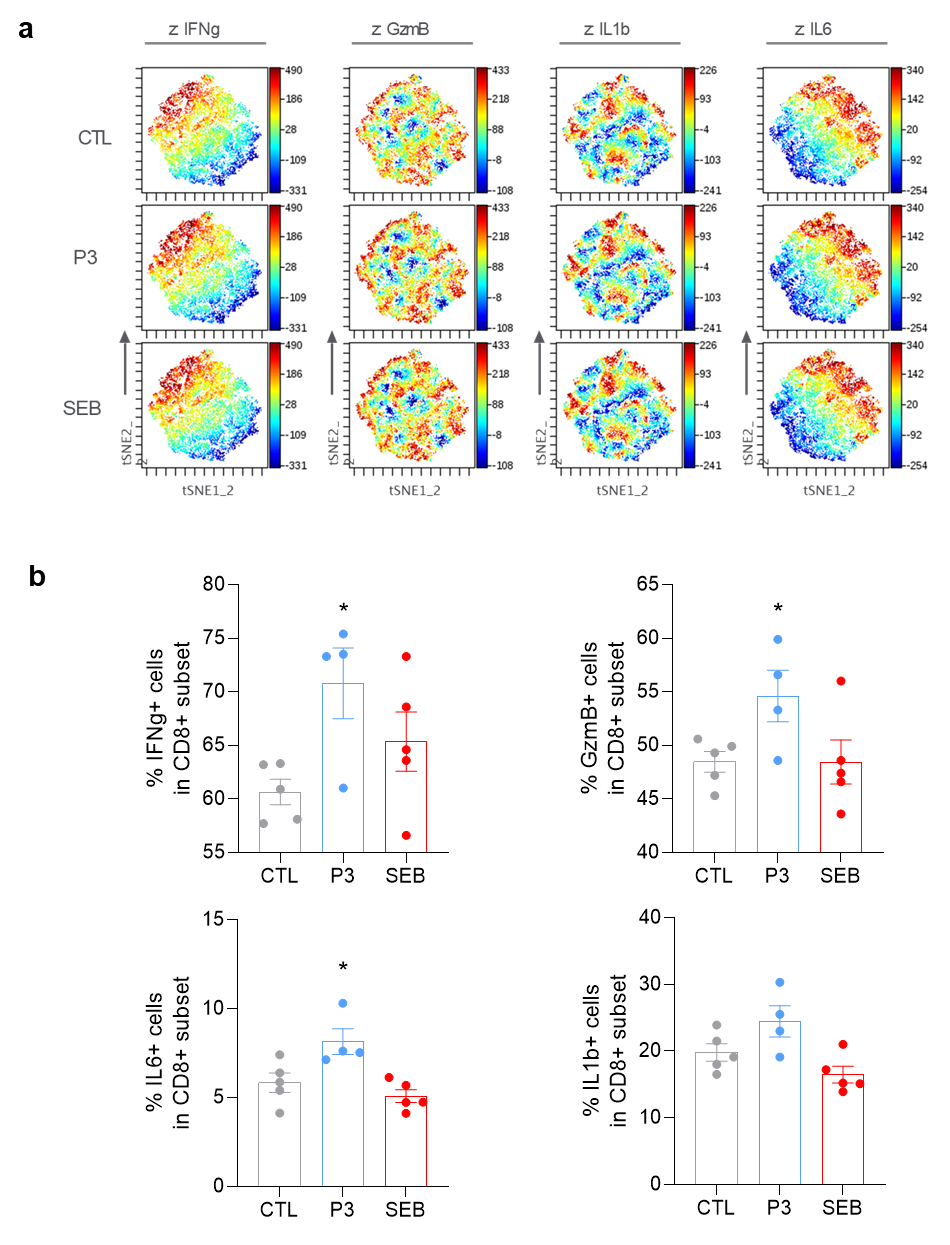

**Supplementary Fig.9. P3 induces a proinflammatory cytokines in CD8+ human T-cells and *in vivo* in mice.** C57BL/6 mice were injected with P3 or SEB once a week for 3 weeks. At the end of experiment blood samples were collected after 1 week of last injection for investigation. viSNE profiles of IFNg, GzmB, IL1b, and IL6 expression level on CD8+ T-cells. The analysis incorporates concatenated data from 5 mice in CTL group, 4 mice in P3-injected group and 5 mice in SEB-injected group. Equal numbers of cells in each group were analyzed. **b** Secretion protein levels of IFNg, GzmB, IL1b, and IL6 cytokines in CD8+ blood T cells of mice immunized with peptide 3 or SEB. The levels of cytokines were determined by using flow cytometry, showing the percentage of CD8+ T cell expressing each cytokine. Data are presented as Mean ± SEM from 5 mice in CTL group, n=4 mice in P3-injected group and n=5 in SEB-injected group; significant at p ≤ 0.05 *.
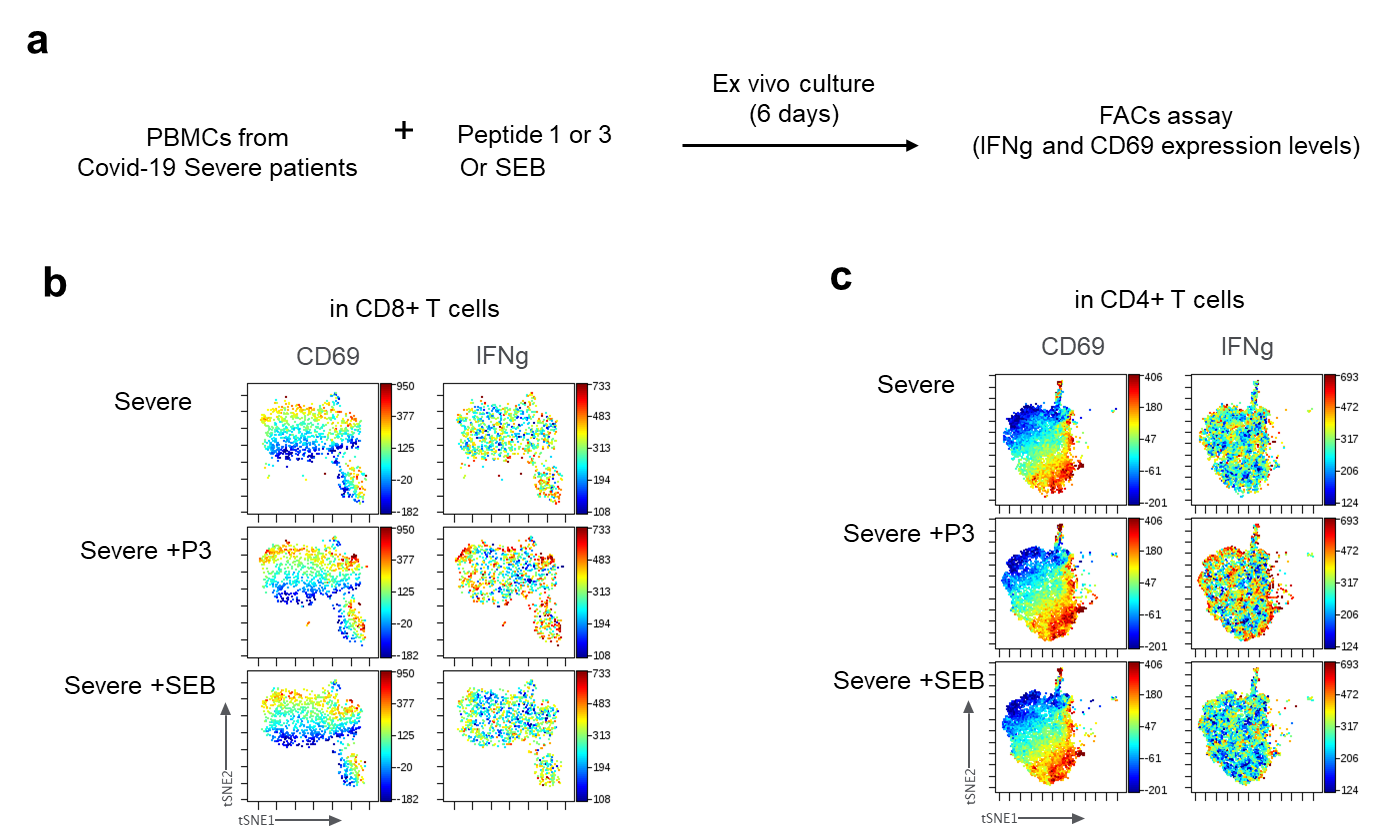

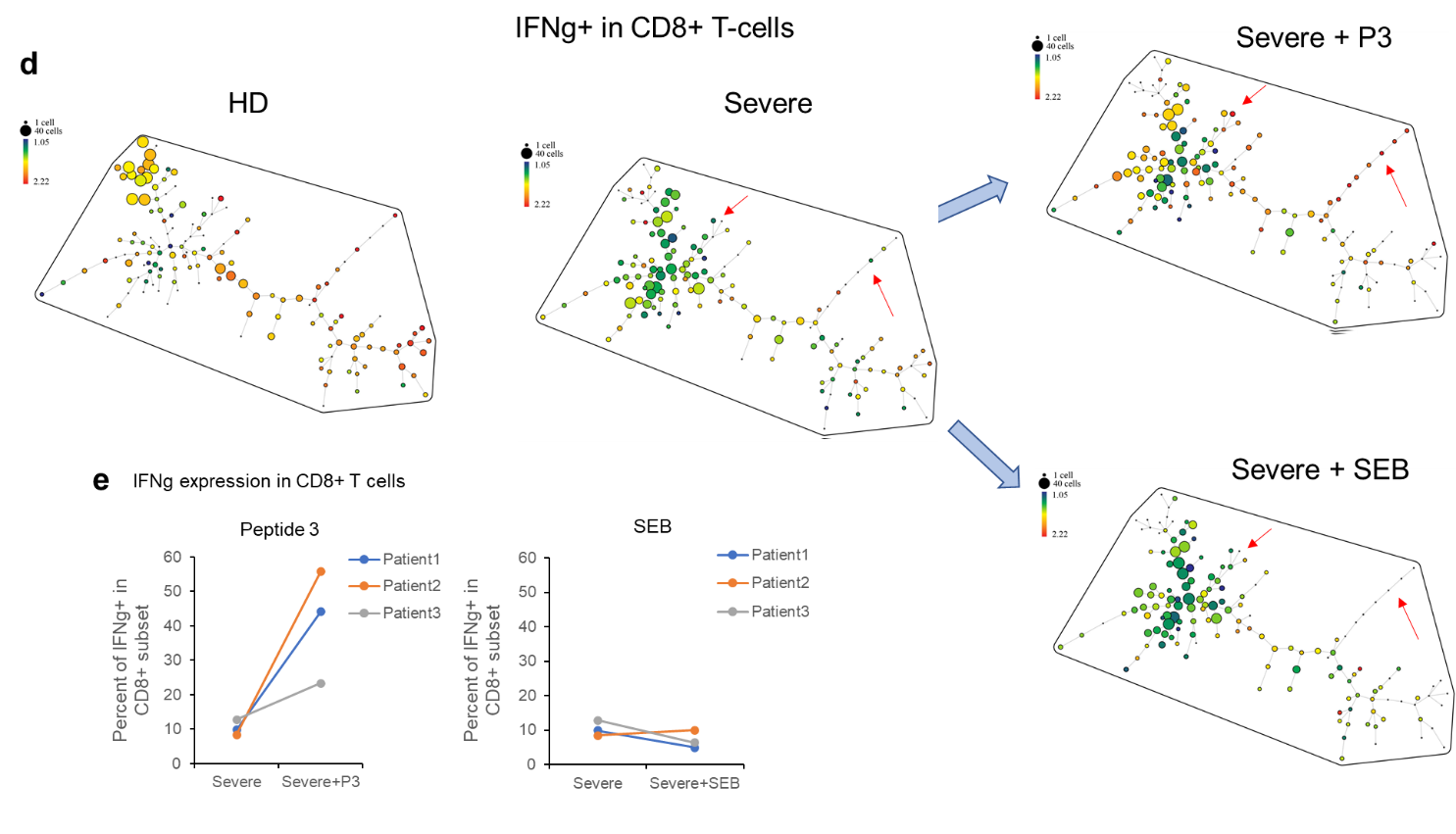

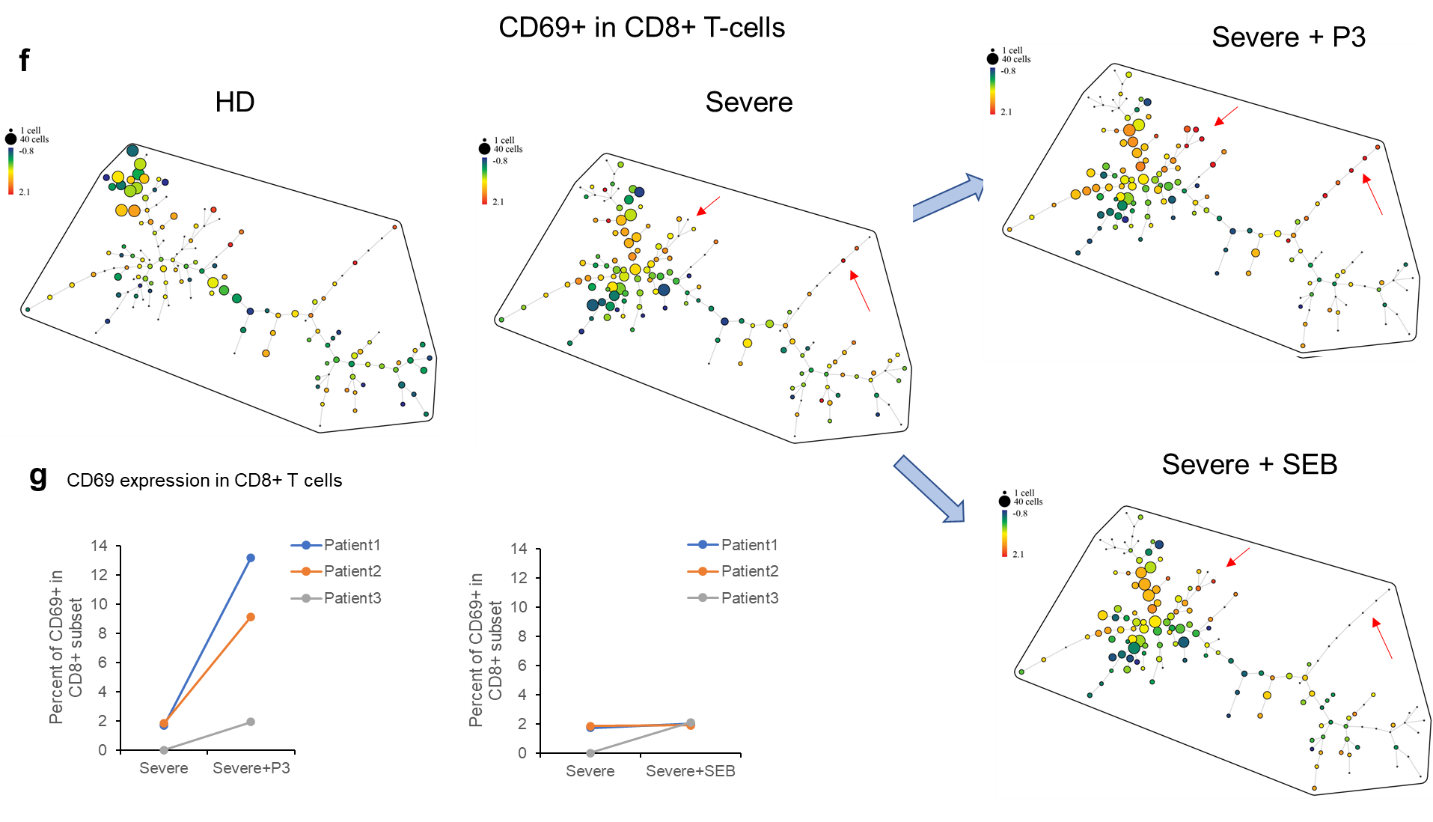

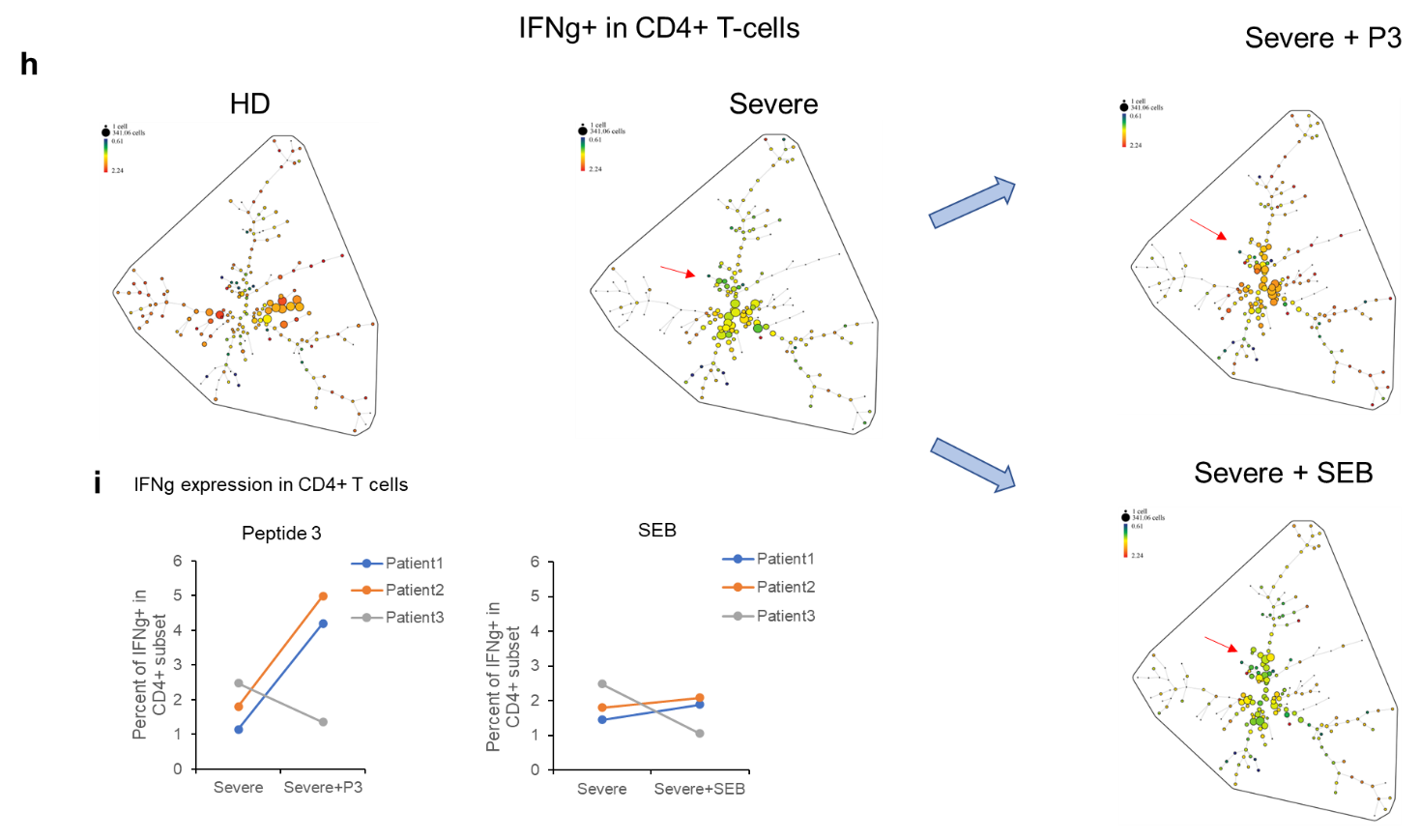

**Supplementary Fig.10. S2 protein peptide rescue CD8+ T-cells of severe Covid-19 patients. a** Flowchart of in vitro experiment showing isolated PBMCs from COVID-19 severe patients were incubated with peptide 3 or SEB peptide. After 6 days, the cells were harvested and stained for flow cytometry analysis. **b** viSNE profiles of CD69 and IFN-g expression on CD8+ T-cells. **c** viSNE profiles of CD69 and IFN-g expression on CD4+ T-cells**. d** SPADE patterns showing the decreasing of IFN-g expression in severe CD8+ T-cells as compare with HD CD8+ T-cells, are recovered with peptide 3 treatment, and no change is seen with SEB peptide. The red narrows show the population with increasing IFN-g expression. **e** Histogram showing the increase of percent of IFN-g+ cells in CD8+ T-cells due to incubation with peptide 3, but not with SEB peptide. The data of PBMCs from COVID-19 severe patients are shown. **f** SPADE patterns showing the decreasing of CD69, activation marker, expression in severe CD8+ T-cells as compare with HD CD8+ T-cells, are recovered with peptide 3 treatment, and no change is seen with SEB peptide. The red narrows show the population with increasing CD69 expression. **g** Histogram showing the increase of percent of CD69+ cells in CD8+ T-cells due to incubation with peptide 3, but not with SEB peptide. The data of PBMCs from COVID-19 severe patients are shown. **h** SPADE patterns showing the decreasing of IFN-g expression in severe CD4+ T-cells as compare with HD CD4+ T-cells, are recovered with peptide 3 treatment, and no change is seen with SEB peptide. The red narrow shows the population with increasing IFN-g expression. **i** Histogram showing the increase of percent of IFN-g+ cells in CD4+ T-cells due to incubation with peptide 3, but not with SEB peptide. The data of PBMCs from COVID-19 severe patients are shown.
